# Evolution of resilience in protein interactomes across the tree of life

**DOI:** 10.1101/454033

**Authors:** Marinka Zitnik, Rok Sosič, Marcus W. Feldman, Jure Leskovec

## Abstract

Phenotype robustness to environmental fluctuations is a common biological phenomenon. Although most phenotypes involve multiple proteins that interact with each other, the basic principles of how such interactome networks respond to environmental unpredictability and change during evolution are largely unknown. Here we study interactomes of 1,840 species across the tree of life involving a total of 8,762,166 protein-protein interactions. Our study focuses on the resilience of interactomes to network failures and finds that interactomes become more resilient during evolution, meaning that interactomes become more robust to network failures over time. In bacteria, we find that a more resilient interactome is in turn associated with the greater ability of the organism to survive in a more complex, variable and competitive environment. We find that at the protein family level, proteins exhibit a coordinated rewiring of interactions over time and that a resilient interactome arises through gradual change of the network topology. Our findings have implications for understanding molecular network structure both in the context of evolution and environment.

---

**T**he enormous diversity of life shows a fundamental ability of organisms to adapt their phenotypes to changing environments (1). Most phenotypes are the result of an interplay of many molecular components that interact with each other and the environment (2–5). The study of life’s diversity has a long history and extensive phylogenetic studies have demonstrated evolution at the DNA sequence level (6–8). While studies based on sequence data alone have demonstrated evolution of genomes, mechanistic insights into how evolution shapes interactions between proteins in an organism remain elusive (9, 10).

DNA sequence information has been used to associate genes with their functions (11), determine properties of ancestral life (12, 13), and understand how the environment affects genomes (14). Despite these advances in understanding DNA sequence evolution, little is known about basic principles that govern the evolution of interactions between proteins. In particular, evolution of DNA and amino acid sequences could lead to pervasive rewiring of protein-protein interactions and create or destroy the ability of the interactions to perform their biological function.

The importance of protein-protein interactions has spurred experimental efforts to map all interactions between proteins in a particular organism, its interactome, namely the complex network of protein-protein interactions in that organism. A large number of high-throughput experiments have reported high-quality interactomes in a number of organisms (15–19). Because interactomes underlie all living organisms, it is critical to understand how these networks change during evolution (20, 21) and elucidate key principles of their structure.

Here, we use protein interactions measured by these large-scale interactome mapping experiments and study the evolutionary dynamics of the interactomes across the tree of life. Our protein interaction dataset contains a total of 8,762,166 physical interactions between 1,450,633 proteins from 1,840 species, encompassing all current protein interaction information at a cross-species scale (SI Appendix, Section 1 and Table S4). We group these interactions by species and represent each species with a separate interactome network, in which nodes indicate a species’ proteins and edges indicate experimentally documented physical interactions, including direct biophysical protein-protein interactions, regulatory protein-DNA interactions, metabolic pathway interactions, and kinase-substrate interactions measured in that species. We integrate into the dataset (22) the evolutionary history of species provided by the tree of life constructed from small subunit ribosomal RNA gene sequence information (12) (SI Appendix, Section 2). Using network science, we study the network organization of each interactome; in particular, its resilience to network failures, a critical factor determining the function of the interactome (23–26). We identify the relationship between the resilience of an interactome and evolution and use this resilience to uncover relationships with natural environments in which organisms live. Although the interactomes are incomplete and biased toward much-studied proteins and model species (SI Appendix, Section 1 and Fig. S7), our analyses give results that are consistent across taxonomic groups, that are not sensitive to network data quality or network size change (SI Appendix, Section 8 and Fig. S8), and indicate that our conclusions will still hold when more protein interaction data become available.

### Significance Statement

The interactome network of protein-protein interactions captures the structure of molecular machinery that underlies organismal complexity. The resilience to network failures is a critical property of the interactome as the breakdown of interactions may lead to cell death or disease. By studying interactomes from 1,840 species across the tree of life, we find that evolution leads to more resilient interactomes, providing evidence for a longstanding hypothesis that interactomes evolve favoring robustness against network failures. We find that a highly resilient interactome has a beneficial impact on the organism’s survival in complex, variable, and competitive habitats. Our findings reveal how interactomes change through evolution and how these changes affect their response to environmental unpredictability.

## Results

### Modeling Resilience of the Interactome

Natural selection has influenced many features of living organisms, both at the level of individual genes (27) and at the level of whole organisms (13). To determine how natural selection influences the structure of interactomes, we study the resilience of in-teractomes to network failures (23, 25, 26). Resilience is a critical property of an interactome as the breakdown of proteins can fundamentally affect the exchange of any biological information between proteins in a cell (Fig. 1a). Network failure could occur through the removal of a protein (*e.g*., by a nonsense mutation) or the disruption of a protein-protein interaction (*e.g*., by environmental factors, such as availability of resources). The removal of even a small number of proteins can completely fragment the interactome and lead to cell death and disease (4, 5) (SI Appendix, Section S5.1 and Table S3). Disruptions of interactions can thus affect the interactome to the extent that its connectivity can be completely lost and the interactome loses its biological function and increases the risk of disease (5).

**Fig. 1.**
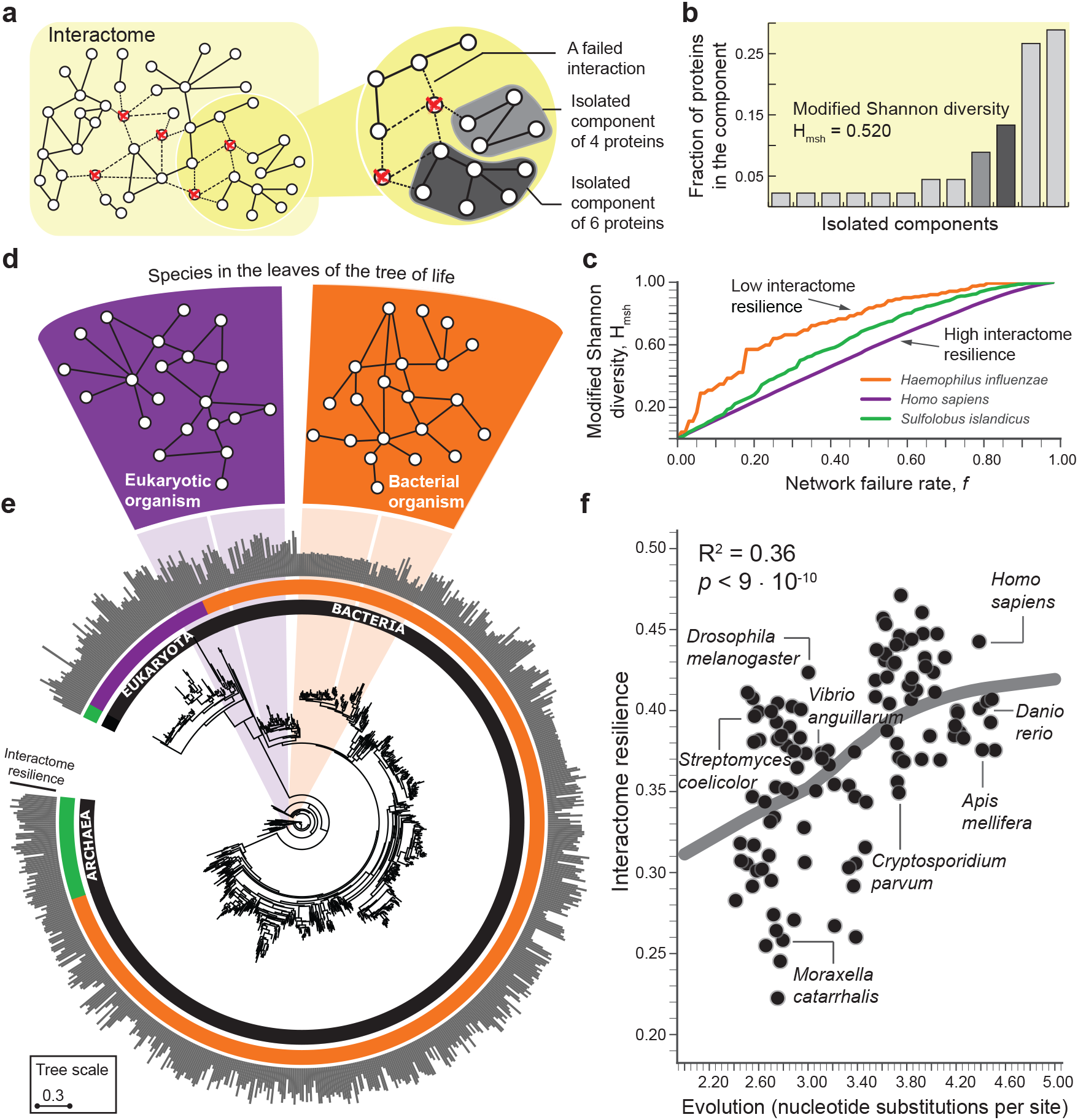
Protein interaction data of 1,840 species consisting of 8,762,166 interactions by 1,450,633 proteins reveal the resilience of interactomes across vast evolutionary distances. (**a**) The interactome of an organism consists of all physical interactions between proteins in the organism. When interactions involving a certain fraction (*f* = 5/45 in this example) of the proteins are removed from the interactome, the interactome fragments into a number of isolated network components. (**b**) Modified Shannon diversity *H*_msh_ (SI Appendix, Section 5) measures how the interactome fragments into isolated components at a given network failure rate *f*. (**c**) The resilience of the interactome integrates modified Shannon diversity *H*_msh_ across all possible failure rates *f* (SI Appendix, Section 5). Resilience value 1 indicates the most resilient interactome, and resilience value 0 indicates a complete loss of the connectivity of the interactome (SI Appendix, Fig. S3). *H. sapiens* (*H. influenzae*) has the most (least) resilient interactome (their resilience is 0.461 and 0.267, respectively) among the three selected organisms. (**d**) A small neighborhood of the interactome in an eukaryotic and a bacterial species. As ancestral species have gone extinct, older interactomes have been lost, and only interactomes of present-day species are available to us. (**e**) Phylogenetic tree showing 1,539 bacteria, 111 archaea, and 190 eukarya (12). Evolution of a species is defined as the total branch length (nucleotide substitutions per site) from the root to the corresponding leaf in the tree (SI Appendix, Section 2). The outside circle of bars shows the interactome resilience of every species. Current protein-protein interaction data might be prone to notable selection and investigative biases (SI Appendix, Section 1); the subsequent plot shows the interactome resilience for 171 species with at least 1,000 publications in the NCBI Pubmed (SI Appendix, Fig. S7). (**f**) Across all species, evolution of a species predicts resilience of the species’ interactome to network failures (LOWESS fit; *R*^2^ = 0.36); more genetic change implies a more resilient interactome. Three species with the most nucleotide substitutions per site (far right on the x-axis) have on average 20.4% more resilient interactome than the three species with the least substitutions (far left on the x-axis).

We formally characterize the resilience of an interactome of a species by measuring how fragmented the interactome becomes when all interactions involving a fraction *f* of the proteins (nodes) are randomly removed from the interactome (Fig. 1a). The resulting isolated network components then determine the interactome fragmentation. A network component is a connected subnetwork of the interactome in which any two nodes can reach each other by a path of edges. The smaller the network component, the fewer nodes can be reached from any given node in the component. To characterize how the interactome fragments into isolated components we use the Shannon diversity index (28–31), which we modify to ensure that the resilience of interactomes with different numbers of proteins can be compared (Fig. 1b and SI Appendix, Section S5.2). In particular, when the interactome *G* is subjected to a network failure rate *f* it is fragmented into a number of isolated components of varying sizes (SI Appendix, Fig. S1). We quantify connectivity of the resulting fragmented interactome *G_f_* by calculating the modified Shannon diversity on the resulting set of isolated components. Let {*C*_ı_, *C_2_,…, C_k_*} be *k* isolated components in *G_f_*. The modified Shannon diversity of *G_f_* is then calculated as the entropy of {*C_1_, C_2_,…,C_k_*} as follows:

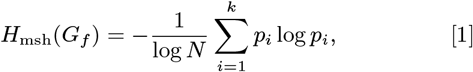

where *N* is the number of proteins in the interactome and *P_i_ = |C_i_|/N* is the proportion of proteins in the interactome *G* that are in component *C_i_*. We can interpret *p_i_* as the probability of seeing a protein from component *C_i_* when sampling one protein from the fragmented interactome *G_f_*. That is, Eq. [1] quantifies the uncertainty in predicting the component identity of a protein that is taken at random from the interactome. Finally, we use the normalization factor 1 */* log *N* because it corrects for differences in the numbers of proteins in the interactomes and ensures that interactomes of different species can be compared. The range of possible *H_msh_* values is between 0 and 1, where these limits correspond, respectively, to a connected interactome in which any two proteins are connected by a path of edges and a completely fragmented interactome in which every protein is its own isolated component. If the fragmented interactome has one large component and only a few small broken-off components, then the modified Shannon diversity is low, providing evidence that the interactome has network structure that is resilient to network failures (23) (SI Appendix, Fig. S2). In contrast, if the interactome breaks into many small components, it becomes fragmented, and its modified Shannon diversity is high (SI Appendix, Fig. S2), indicating that the interactome is not resilient to network failures.

To fully characterize the interactome resilience of a species we measure fragmentation of the species’ interactome across all possible network failure rates (SI Appendix, Fig. S3). Consider the interactome of the pathogenic bacterium *H. influenzae* and the interactome of humans, which have different resilience (Fig. 1c). In the *H. influenzae* interactome, on removing small fractions of all nodes many network components of varying sizes appear, producing a quickly increasing Shannon diversity. In contrast, the human interactome fragments into a few small components and one large component whose size slowly decreases as small components break off, resulting in Shannon diversity that increases linearly with the network failure rate (Fig. 1c). Thus, unlike the fragmentation of the *H. influenzae* interactome, the human interactome stays together as a large component for very high network failure rates, providing evidence for the topological stability of the interactome. In general, the calculation of modified Shannon diversity over all possible network failure rates *f* yields a monotonically increasing function that reaches its minimum value of 0 at *f* = 0 (*i.e*., a connected interactome) and its maximum value of 1 at *f* =1 (*i.e*., a completely fragmented interactome) as the interactome becomes increasingly fragmented with increasing network failure rate *f* (SI Appendix, Section 5.3 and Fig. S3). We therefore define resilience of interactome *G* as one minus the area under the curve defined by that function:

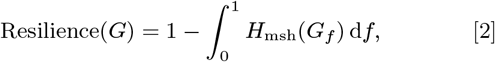

which takes values between 0 and 1; a higher value indicates a more resilient interactome.

### Resilience of Interactomes Throughout Evolution

We characterize systematically the resilience of interactomes for all species in the dataset (Fig. 1d and SI Appendix, Table S5). We find that species display varying degrees of interactome resilience to network failures (Fig. 1e). At a global, cross-species scale, we find that a greater amount of genetic change is associated with a more resilient interactome structure (LOWESS fit; *R*^2^ = 0.36; Fig. 1f), and this association remains strong even after statistical adjustments for the influence of many other variables (SI Appendix, Section 8). The more genetic change a species has undergone, the more resilient is its interactome. The evolution a species, which is defined by the total branch length from the root to the leaf taxon representing that species in the tree of life (12), thus predicts resilience of the species’ interactome, providing empirical evidence that interactome resilience is an evolvable property of organisms (26). This finding also suggests that the structure of present-day interactomes reflects their history or that interactomes must have a certain structure because that structure is well suited to the network’s biological function. From an evolutionary standpoint, this finding points in the direction of topologically stable interactomes, which suggests that evolutionary forces may shape protein interaction networks in such a way that their large-scale connectivity, *i.e*., the network’s biological function, remains largely unaffected by small network failures as long the failures are random.

We also find that species from the same taxonomic domain have more similar interactome resilience than species from different domains (*p* =6 · 10^-11^ for bacteria against eukaryotes; see SI Appendix, Fig. S10 for comparisons between other taxonomic groups). Furthermore, the degree of interactome resilience is significantly higher than expected by chance alone (SI Appendix, Fig. S9); that is, in a similar random network of identical size and degree distribution (*p* = 5·10^-12^), indicating that naturally occurring interactomes have higher resilience than their random counterparts. These findings are independent of genomic attributes of the species, such as genome size and the number of protein-coding genes, and are not direct effects of network size, the number of interactions in each species, broad-tailed degree distributions (23), or the presence of hubs in the interactome networks (SI Appendix, Fig. S8 and Table S1). Furthermore, these findings are consistent across a variety of assays that are used to measure the interactome (SI Appendix, Table S2).

### Relationship between Interactome Resilience and Ecology

We next ask if there is a relationship between species’ interactome resilience and aspects of species’ ecology (SI Appendix, Section 4). We examine the relationship between interactome resilience and the fraction of regulatory genes and find that bacteria with more resilient interactomes have significantly more regulatory genes in their genomes (*R*^2^ = 0.32; Fig. 2a). Bacteria with highly resilient interactomes can also survive in more diverse and competitive environments, as revealed by exceptionally strong associationss between the resilience and the level of co-habitation and the environmental scope (Fig. 2b-c). Furthermore, using a categorization of bacteria into five groups based on their natural environments [NCBI classification for bacterial lifestyle (terrestrial, multiple, host cell, aquatic, specialized) ordered by the complexity of each category (32)], we find that terrestrial bacteria living in the most complex ecological habitats have the most resilient interactomes (*p* = 7 · 10^-3^; Mann-Whitney U test), and that host-associated bacteria have the least resilient interactomes (*p* = 4 · 10^-6^; Fig. 2d). Our analysis further reveals that interactome resilience is indicative of oxygen dependence; the most resilient interactomes are those of aerobic bacteria (*p* = 8 · 10^-4^), followed by facultative and then anaerobic bacteria, which do not require oxygen for growth (Fig. 2d).

**Fig. 2.**
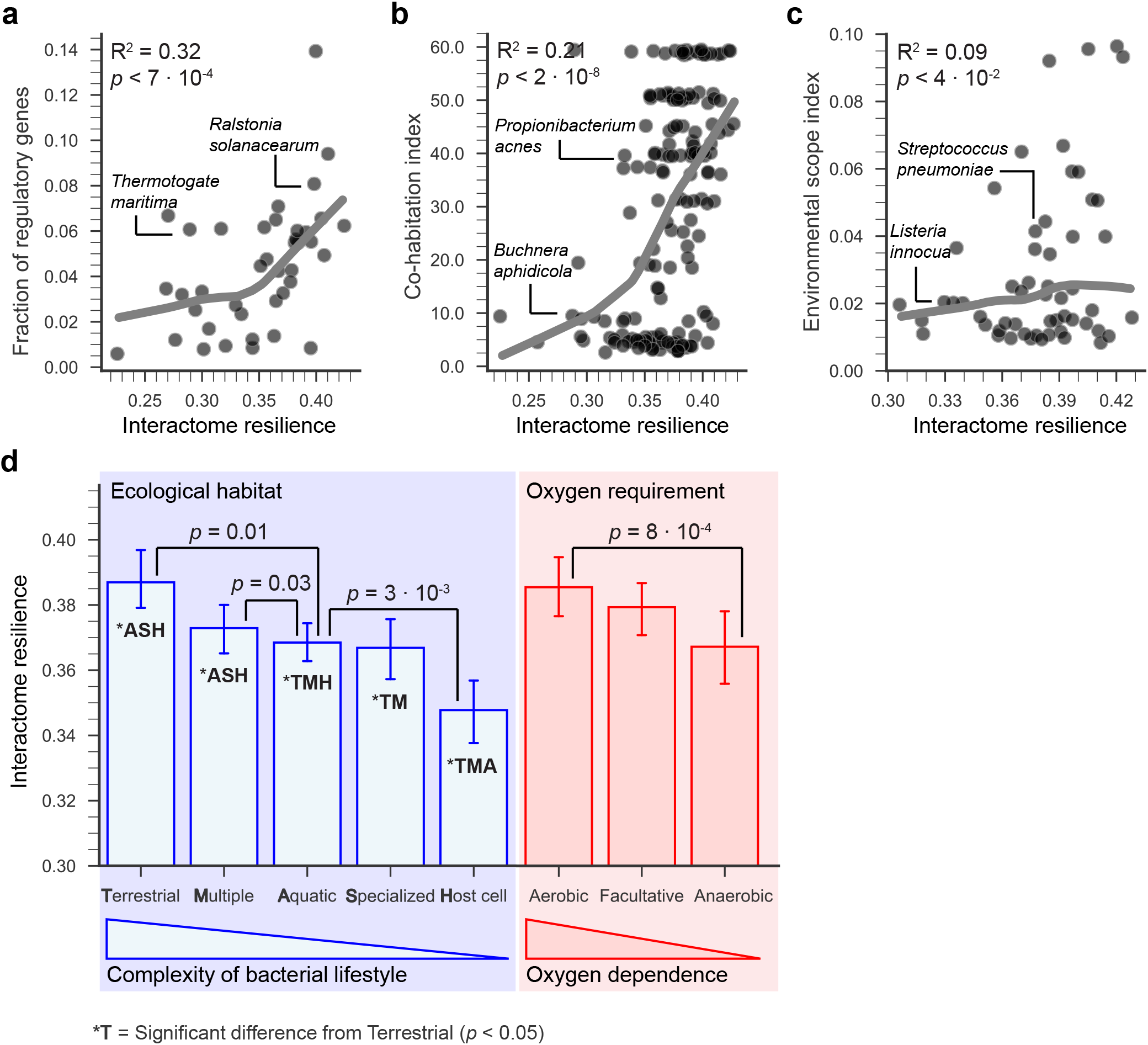
Bacteria with more resilient interactomes survive in more complex, variable and competitive environments. We use ecological information for 287 bacterial species (32) to examine the relationship between species' interactome resilience and their ecology (SI Appendix, Section 4). (**a**) Interactome resilience positively correlates with the fraction of regulatory genes in bacteria, an established indicator of environmental variability of species' habitats (32) (*R*^2^ = 0.32). (b and c) For environmental viability of a species, we use a co-habitation index that records how many organisms populate each environment in which the species is viable (*i.e*., the level of competition in each viable environment), and an environmental scope index that records a fraction of the environments in which the species is viable (i.e., species' environmental diversity) (32). The resilience of the interactome positively correlates with the level of co-habitation encountered by bacteria (*R*^2^ = 0.21), and bacteria with resilient interactomes tend to thrive in highly diverse environments (*R*^2^ = 0.09). (**d**) Terrestrial bacteria have the most resilient interactomes (*p = 7* · 10^-3^), and host-associated bacteria have the least resilient interactomes (*p* = 4 · 10^-6^). In bacteria, interactome resilience is indicative of oxygen dependence. Aerobic bacteria have the most resilient interactomes (*p* = 8 · 10^-4^), followed by facultative and the anaerobic bacteria. Error bars indicate 95% bootstrap confidence interval; *p* values denote the significance of the difference of the means according to a Mann-Whitney U test.

These relationships suggest that molecular mechanisms that render a species’ interactome more resilient might also allow it to cope better with environmental challenges. In the network context, high interactome resilience suggests that proteins can interact with each other even in the face of high protein failure rate. High interactome resilience indicates that a species has a robust interactome, in which many mutations represent network failures that are neutral in a given environment, have no phenotypic effect on the network’s function and are thus invisible to natural selection (26). However, neutral mutations may not remain neutral indefinitely, and a once-neutral mutation may have phenotypic effects in a changed environment and be important for evolutionary innovation (25). Although a large number of mutations in a resilient interactome might not change the network’s primary function, they might alter other network features, which can drive future adaptations as the environment of the species changes (33). Changes that are neutral concerning one aspect of the network’s function could lead to innovation in other aspects, suggesting that a resilient interactome can harbor a vast reservoir of neutral mutations.

### Structural Changes of Protein Network Neighborhoods

A resilient interactome may arise through changes in the network structure of individual proteins over time (Fig. 3a). To investigate such changes in local protein neighborhoods, we decompose species interactomes into local protein networks, using a 2-hop subnetwork centered around each protein in a given species as a local representation of a protein’s direct and nearby interactions in the species’ interactome (SI Appendix, Section 6.1 and Fig. S4). We obtain 81,673 protein network neighborhoods and then use orthologous relationships between proteins to group them into 2,224 protein families, with an average of 38 protein neighborhoods originating from 12 species in each family (SI Appendix, Section 3). Each family represents a group of orthologous proteins that share a common ancestral protein (Fig. 3b).

**Fig. 3.**
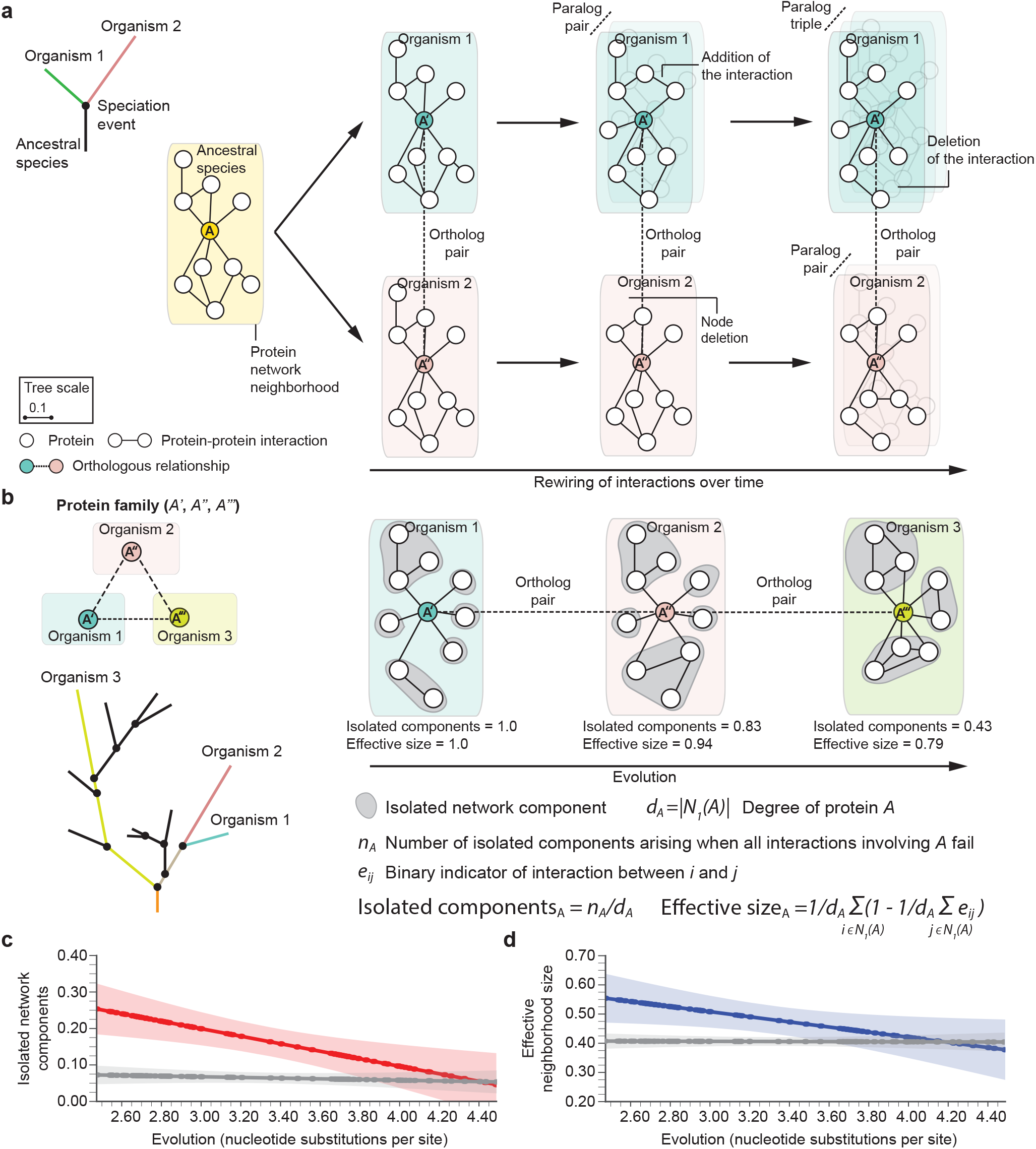
Evolution mitigates local network structural changes in protein interactomes. (**a**) A hypothetical phylogenetic tree illustrates a speciation event that gives rise to two lineages according to the speciation-divergence model (11) and leads to present-day organisms ‘1’ (green) and ‘2’ (pink). In this example, a single ancestral protein *A* that was present in the ancestral species gives rise to proteins *A’* and *A’’* upon speciation; *A’* and *A’’* form an orthologous protein pair. As the two newly arising species diverge and protein sequences evolve, protein network neighborhoods (SI Appendix, Fig. S4) in their interactomes can rewire independently overtime. Shown are also in-paralogs, proteins which arise through gene duplication events in species ‘1’ and ‘2’ after speciation. (**b**) A hypothetical protein family with three protein members (*A’, A’’, A’’’*), each from a different organism. In the phylogenetic tree, organism ‘1’ is located at the tip of the lineage with the shortest branch length, whereas organism ‘3’ is in the lineage with the longest branch length in the tree. We represent the protein family by a sequence of orthologous proteins ordered by the branch length of proteins’ originating species (SI Appendix, Section 3). We then characterize the network neighborhood of each protein in the family by calculating two network metrics (SI Appendix, Fig. S5). Isolated components are given by the degree-adjusted number of connected components in the neighborhood that arise when the central protein is removed from the interactome (gray) (SI Appendix, Section 6). The neighborhood size down-weighted by the redundancy of local interactions gives the effective size of the neighborhood (SI Appendix, Section 6). (c and d) The number of isolated network components and the effective size of protein neighborhoods both decrease with evolution (*p* = 3 · 10^-8^ and *p* = 0.03, respectively; Spearman’s *ρ* rank correlation), suggesting that local interaction neighborhoods rewire via a coordinated evolutionary mechanism. Lines in c and d show the LOWESS fit of median-aggregated network metric values for 81,673 proteins from 2,224 protein families; color bands indicate 95% confidence band for the LOWESS fit; gray lines show random expectation.

By examining protein families, we find that the number of isolated network components in protein network neighborhoods and the effective size of the neighborhoods (Fig. 3b; SI Appendix, Section 6.2) both decrease with evolution (*p* = 3 · 10^-8^ and *p* = 0.03, respectively; Fig. 3c-d), indicating that protein neighborhoods become more connected during evolution. These structural changes in the neighborhoods suggest a molecular network model of evolution (Fig. 3b): For orthologous proteins in two species, as the evolutionary distance between the species increases, the proteins’ local network neighborhoods become increasingly different and the neighborhood becomes more interconnected in the species that has undergone more genetic change.

### Network Rewiring of Protein-Protein Interactions

To study evolutionary mechanisms of structural changes in the interactomes, we investigate network motifs (34, 35). We first identify orthologous protein pairs from evolutionarily close species (SI Appendix, Section 3), resulting in 2,485,564 protein pairs, which we then use to calculate interaction rewiring rates (IRR) for selected network motifs (Fig. 4a). We calculate the number of times each motif appears in each protein neighborhood and derive the IRR by comparing the motif occurrences between the interactomes of the older and the younger species of each protein pair (SI Appendix, Section 7.1). We find strong statistical evidence that network motifs rewire during evolution (*p* < 10^-33^ for all network motifs; Fig. 4b), suggesting that rewiring of interactions is an important mechanism for the evolution of interactomes. For example, proteins in evolutionarily older species on average participate in a factor of 0.861 fewer protein-protein interactions compared to proteins in evolutionarily younger species (IRR = -0.215; Fig. 4b). This significant negative correlation between each protein’s number of interactions and the protein’s evolutionary age confirms earlier studies in *S. cerevisiae* (36). We also find that square motifs of interactions become more common in protein neighborhoods during evolution (IRR = 0.016; Fig. 4b). A range of biological evidence (18, 37, 38) supports this positive rate of change in the number of square motifs: From a structural perspective (38), protein-protein interactions often require complementary interfaces; hence two proteins with similar interfaces share many of their neighbors. However, they might not interact directly with each other, which manifests in the interactome as a square motif of interactions (see SI Appendix, Fig. S6 for an illustration of interaction interfaces recognizing the binding sites in proteins). Evolutionary arguments following gene duplication (18) reach the same conclusion; proteins with multiple shared interaction partners are likely to share even more partners and thereby produce new square motifs of interactions. To test the predictive power of our motif-based model of structural network changes, we estimate the size of the whole human interactome by extrapolating the *S. cerevisiae* interactome, using interaction rewiring rates from Fig. 4b (SI Appendix, Section 7.3). Assuming one splice isoform per gene, we predict the number of interactions in humans to be ~160,000. This prediction is in surprisingly good agreement with three previous estimates of the size of the human interactome, which range from 150,000 to 370,000 interactions (15, 17, 39) and have proved crucial in establishing the complexity of the human interactome (19).

**Fig. 4.**
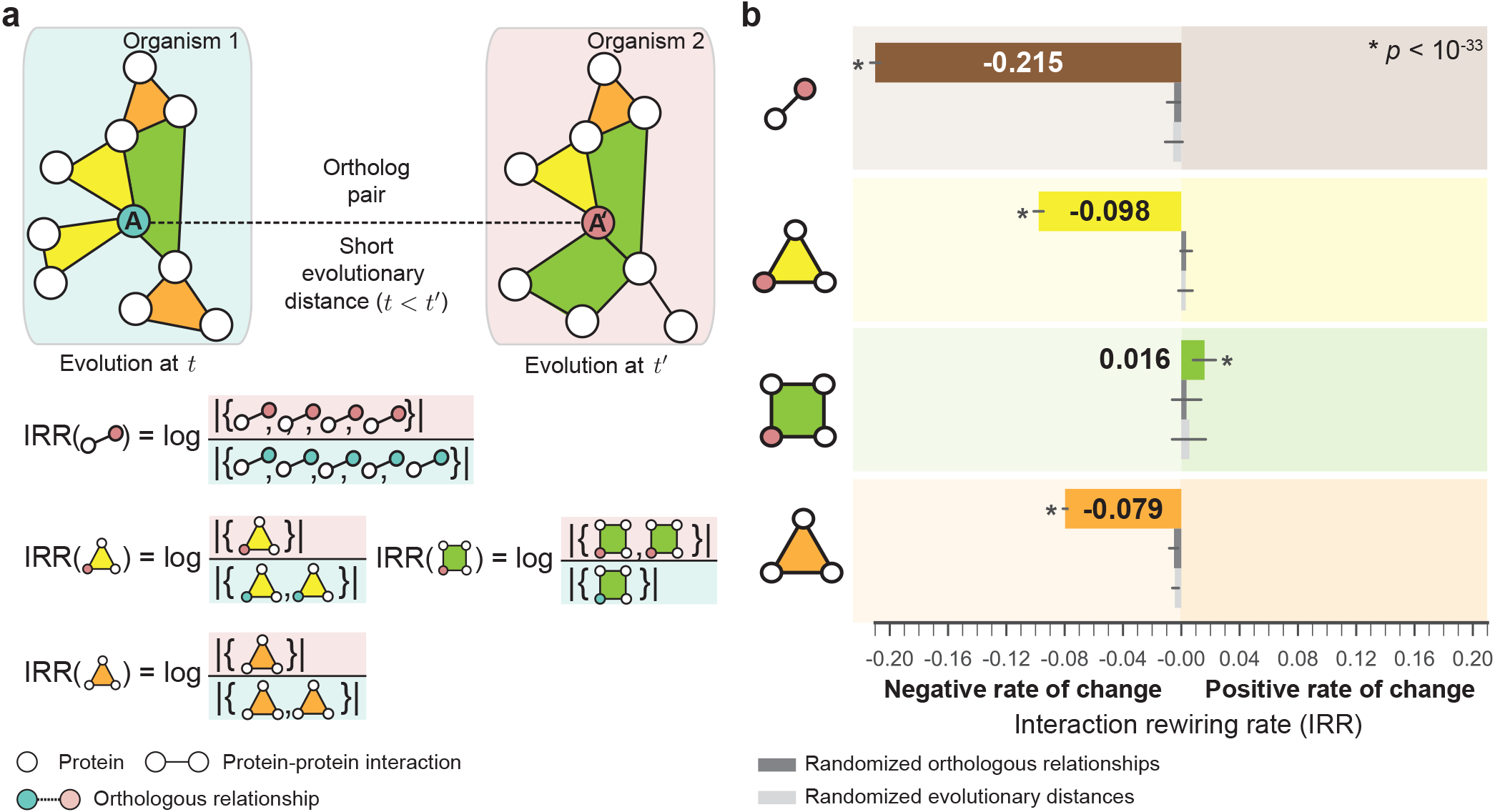
The rewiring rate of interactions in local protein neighborhoods varies with the topology of network motifs. (**a**) Interaction rewiring rate (IRR) measures the fold change between the probability of observing a particular network motif in the network neighborhood of protein *A’* relative to the probability of observing the same motif in the neighborhood of an evolutionarily younger orthologous protein A. A positive (negative) rate of change indicates the motif becomes more (less) common over time (SI Appendix, Section 7). Shown are the rewiring rates for interactions (*i.e*., edges; the number of interactors of *A’* versus A), triangle motifs touching the orthologous protein (yellow), square motifs touching the orthologous protein (green), and triangle motifs in the protein network neighborhood (orange). (**b**) Square motifs become more common in protein neighborhoods during evolution (*p* < 10^-33^), which is supported by a range of biological evidence (18, 37, 38). However, triangle motifs become less common over time (*p* < 10^-33^ for both types of triangle motifs). Gray bars indicate random expectation (SI Appendix, Section 7), either for random orthologous relationships (dark gray) or for random evolutionary distances (light gray); error bars indicate 95% bootstrap confidence interval; *p* values denote the significance of the difference of IRR distributions using a two-sample Kolmogorov-Smirnov test.

## Discussion

Our analyses reveal how protein-protein interaction networks change through evolution and how changes in these networks affect phenotypes and organismal response to environmental complexity. This systematic investigation of protein-protein interaction networks from an evolutionary perspective was enabled by a dataset of interactomes, consisting of protein-protein interaction networks from 1,840 species. To date, most evolutionary analyses of biological networks have focused on a small number of organisms with high-coverage protein-protein interaction data, such as *S. cerevisiae, M. musculus*, and humans. This is because interactomes mapped by unbiased tests of all possible pairwise combinations of proteins on the same platform remain scarce, an important limitation of the present study. Furthermore, experimentally documented protein interactions are currently subject to a high number of false positives and negatives. As more protein interaction data are collected, and more genomes become available, the generalizability of our findings can be further evaluated. However, our results are consistent across both different subsets of protein interaction data (SI Appendix, Table S2) and different phylogenetic lineages (SI Appendix, Fig. S10) and are not explained by many possible genomic and network confounders (SI Appendix, Section 8, Fig. S8, and Table S1), thus providing confidence that our key findings cannot be attributed to biases in the datasets.

Interactome resilience is an important aspect of our study. The resilience measures fragmentation of the interactome into isolated components and thus represents a global measure of the interactome’s topological stability. Beyond fragmentation, there are other possible modifications of the interactome that could alter the network’s biological function without necessarily disconnecting the network (40–42). As more detailed information about functions of individual proteins in the interactome (43), as well as dynamic protein-expression data (44), becomes available, our measure of interactome resilience could be adapted to give a more complex definition of resiliency, which might yield more detailed evolutionary predictions. Additionally, information on how protein-protein interactions change dynamically both in time and space (45–47) might reveal how topological stability of the interactome depends on large-scale interactome connectivity as well as on the interactome’s dynamic properties (40).

Our study presents a new paradigm for evolutionary studies by demonstrating that interactomes reveal fundamental structural principles of molecular networks. Our findings highlight evolution as an important predictor of structural network change and show that evolution of a species predicts resilience of the species’ interactome to protein failures. The findings offer quantitative evidence for the biological proposition that an organism that has undergone more genetic change has a more resilient interactome, which, in turn, is associated with the greater ability of the organism to survive in a more complex, variable or competitive environment. Our findings can also help clarify the mechanisms of how interactomes change during evolution, why currently observed network structures exist and how they may change in the future, and facilitate the extrapolation of functional information from experimentally characterized proteins to their orthologous proteins in poorly studied organisms.

## Materials and Methods

Detailed description of data, statistical methodology, and additional analyses are provided in SI Appendix.

## Code and Data Availability

Software implementation of statistical methodology is publicly available at http://snap.stanford.edu/tree-of-life. All data used in the paper, including the processed interactomes, are shared with the community and available from http://snap.stanford.edu/tree-of-life.

## ACKNOWLEDGMENTS

For helpful discussion and comments on the manuscript, we thank Aaron Goodman, Hunter B. Fraser, Alice Y. Ting, Emma Pierson, Michael An, and Pratyaksh Sharma. M.Z., R.S., and J.L. were supported in part by NSF, NIH, DARPA, Boeing, Stanford Data Science Initiative, and Chan Zuckerberg Biohub. M.W.F. was supported in part by the Stanford Center for Computational, Evolutionary and Human Genomics and a grant from the John Templeton Foundation.

