## Supplementary material for "Evolution of resilience in protein interactomes across the tree of life": SI Appendix

This PDF file includes:

Supplementary text (**SI Appendix**)

Supplementary figures S1 to S10

Supplementary tables S1 to S5

Supplementary references

#### Supplementary Text (SI Appendix)

**S1 Protein-protein interaction dataset** **S4**
S1.1 Protein-protein interaction network data . . . . . S4
S1.2 Biases in the protein-protein interaction dataset . . . . . S7

**S2 The tree of life dataset** **S8**
S2.1 Phylogenetic tree of species . . . . . S9
S2.2 The NCBI Taxonomy database . . . . . S10

**S3 Information on clusters of orthologous genes and protein families** **S10**

**S4 Information on natural environments of species** **S11**

**S5 Additional information on interactome resilience** **S13**
S5.1 Motivation and overview of the approach . . . . . S13
S5.2 Modified Shannon diversity . . . . . S15
S5.3 Interactome resilience . . . . . S17
S5.4 Removal of nodes representing essential protein-coding genes . . . . . S18

**S6 Additional information on analysis of protein network neighborhoods** **S19**
S6.1 Protein network neighborhoods . . . . . S19
S6.2 Analyses of protein network neighborhoods . . . . . S20

**S7 Additional information on analysis of interactome networks** **S22**
S7.1 Protein-protein interaction rewiring rates (IRR) . . . . . S22
S7.2 Interactome network null models . . . . . S24
S7.3 Estimating the size of the whole human interactome . . . . . S25

**S8 Additional analyses on possible confounding factors** **S27**
S8.1 Confounding factors and partial correlation analyses . . . . . S28
S8.2 Comparison with unbiased datasets . . . . . S30

**Supplementary references** **S53**

#### Supplementary Figures

S1 Characterizing fragmentation of the interactome into isolated components upon node removal S32
S2 Quantifying fragmentation of the interactome using modified Shannon diversity . . . . . S33
S3 Interactome resilience . . . . . S34
S4 Protein network neighborhoods in the interactome . . . . . S35

S5 Characterization of protein network neighborhoods . . . S36
S6 Square network motifs of protein-protein interactions . . . S37
S7 Publication bias towards model organisms and highly studied species . . . S38
S8 Causal model for alternative hypotheses to explain the relationship between evolution and interactome resilience . . . S39
S9 Relationship between evolution and interactome resilience under random expectation . . . S40
S10 Interactome resilience for species from the same taxonomic groups . . . S41

#### Supplementary Tables

S1 Analysis of possible confounding factors for interactome resilience . . . S42
S2 Quality-controlled analysis of interactome data generated by yeast two-hybrid assays . . . S43
S3 Resilience of species' interactomes to network failure of essential protein-coding genes . . . S44
S4 Summary of dataset statistics for species and their genomes . . . S45
S5 Summary of interactome resilience and dataset statistics for species and interactomes . . . S48

#### Supplementary Text

In this Supplementary Text, we present a detailed discussion of the datasets and their analysis. The text is organized as follows. We first describe in detail all biological data we used: the protein-protein interaction dataset (Section S1), phylogenetic information (Section S2), taxonomic information (Section S3), and ecological information about species (Section S4). We then derive our interactome resilience approach introduced in the main text, provide additional explanations and examples, and discuss interactome resilience in more detail than in the main text (Section S5). We also describe our analyses of protein network neighborhoods (Section S6) and the different statistical tests and controls (Section S7). Finally, we present our analyses of the impact of data biases and false positives on our main results (Section S8).

We then present supplementary figures (Figure S1-Figure S10) and tables (Table S1-Table S5).

#### S1 Protein-protein interaction dataset

In this section, we describe how we compile the protein-protein interaction dataset.

##### S1.1 Protein-protein interaction network data

In building the interactomes, we rely only on physical protein-protein interactions that are experimentally supported or manually curated, hence we do not include interactions extracted from gene expression data, evolutionary considerations, and computational predictions. In order to obtain the interactomes as complete as currently feasible, we combine several kinds of physical protein-protein interactions (*1*), including regulatory interactions, binary interactions derived from yeast-two-hybrid high-throughput datasets, metabolic enzyme-coupled interactions, protein complexes, kinase-substrate pairs, and signaling interactions.

**Compilation of the protein-protein interaction dataset.** We collect and reassess experimental and curated data on protein-protein interactions, known pathways, and protein complexes from the raw STRING (Search Tool for the Retrieval of Interacting Genes/Proteins) database (http://string-db.org, obtained "License of STRING for Scientific Purposes" from the European Molecular Biology Laboratory (EMBL); March 16, 2016) (*2, 3*). Broadly, the STRING database integrates information on protein-protein interactions by consolidating known and predicted protein-protein association data for a large number of organisms/species. The associations in STRING include physical (direct) interactions, as well as functional (indirect) interactions, as long as both are specific and biologically meaningful. In this study, however, we specifically focus on physical interactions and thus we exclude functional (indirect) associations from the analysis. We combine the following protein-protein interaction data:

(*a*) *Experimentally supported interactions:* Interactions derived from experiments in the laboratory, including biochemical, biophysical, and genetic assays. Data is populated mainly from the primary protein-protein interaction databases organized in The International Molecular Exchange Consortium (IMEx) consortium (*4*) and The Biological General Repository for Interaction Datasets (BioGRID) (*5*).

(*b*) *Human expert-curated interactions:* Interactions that have been asserted by human expert curators. Data is populated mainly based on known pathways and protein complexes from curated databases. Included are: regulatory interactions from the TRANScription FACtor (TRANSFAC) database (*6*), which lists interactions derived from the presence of a transcription factor binding site in the promoter region of a certain gene; metabolic enzyme-coupled interactions from the Kyoto Encyclopedia of Genes and Genomes (KEGG) database (*7*), which lists interactions derived from coupled enzymes that share adjacent reactions; and protein complexes from the Comprehensive Resource of Mammalian Protein Complexes (CORUM) database (*8*), which lists protein complexes consisting of multiple gene products.

The union of all interactions obtained from (*a*)-(*b*) yields the protein-protein interaction dataset used in this study. The dataset contains 8,762,166 protein-protein interactions defined on 1,450,633 proteins that span 1,840 distinct species (1,539 bacteria, 111 archaea, and 190 eukarya). This dataset is used to construct the interactomes as described in the following paragraphs.

The protein-protein interaction dataset has two appealing features:

($a$) It is a *quality-controlled* dataset as it only includes protein-protein interactions that are supported by either experiments or curated databases rather than computational predictions. More specifically, we discard information that does not represent a strong indication for physical protein-protein interactions, such as information derived from: (i) systematic co-expression analysis (*e.g.*, pairs of proteins that are consistently similar in their expression), (ii) shared signals across genomes (*e.g.*, pairs of proteins that are observed in each other's genome neighborhood such as in the case of conserved and co-transcribed operons), and (iii) automated text-mining of the scientific literature (*e.g.*, pairs of proteins that are frequently mentioned together in the same paper, abstract or even sentence).

($b$) It is a *species-specific* dataset as it only includes interactions that were specifically measured in species. This means the dataset does not include computationally predicted protein-protein interactions generated by techniques that transfer information between species using gene orthology (*e.g.*, (*9*)). More specifically, we discard information on interactions obtained by: (i) computational transfer of interactions between organisms based on gene orthology (*e.g.*, pairs of proteins that have highly similar phylogenetic distributions of orthologs, *i.e.*, if their orthologs tend to be observed as 'present' or 'absent' in the same subsets of organisms), and (ii) computational transfer of interactions between closely related organisms (*e.g.*, pairs of proteins for which there is at least one organism where their respective orthologs have fused into a single, protein-coding gene).

Note that the protein-protein interaction dataset is available through http://snap.stanford.edu/tree-of-life.

**Construction of interactomes.** We take the protein-protein interaction dataset and use it to construct protein-protein interaction networks, called *interactomes* (*1, 10–12*), for a variety of species. In particular, we represent each species $s$ by its interactome, a network $G^{(s)} = (V^{(s)}, E^{(s)})$ in which nodes $V^{(s)}$ represent proteins (protein-coding genes) and edges $E^{(s)}$ represent protein-protein interactions specifically documented in that species. Following earlier literature on the analysis of interactome data (*e.g.*, (*1, 10–12*)), we treat the interactomes as undirected and unweighted (binary) networks.

##### S1.2 Biases in the protein-protein interaction dataset

Currently available protein-protein interaction information is highly biased and only covers a relatively small portion of the proteome, even for the highly studied model organisms and human (*11*). In this study we consider and address two types of data biases and show that our key findings cannot be attributed to these data biases:

(*a*) *Inter-species data bias:* Currently available interactomes vary considerably across different species in how well they recapitulate physical relationships between proteins. This variability comes from the fact that certain species represent major model organisms,[1] which have been widely studied, usually because they are easy to maintain and breed in a laboratory setting and have particular experimental advantages. As a result, major model organisms can have interactomes that are more extensively documented with many characterized protein-protein interactions; however, little can be known about interactomes of organisms that are less widely used in biological research.

(*b*) *Intra-species data bias:* The situation is further complicated by the uneven quality and investigative biases involving experimental interactome mapping pipelines (*e.g.*, a bias in a species towards studying interactions involving proteins encoded by genes that are highly expressed in certain cell lines or associated with certain phenotypes/diseases (*11*)). In particular, this variability means that current interactomes are prone to selection and investigative biases, such as those related to the selection of proteins and the interaction density (number of interactions/edges present in the interactome) (see Figure S8). This variability could potentially indicate that selection and investigative biases, and not fundamental biological properties, underlie the network structure of interactomes.

Next, we describe the analyses performed to address the inter-species data bias.

[1]Model organisms are non-human species that are used in the laboratory to help scientists understand biological processes. Ten most popular model organisms in the U.S. according to NIH: https://publications.nigms.nih.gov/thenewgenetics/poster.pdf.

**Addressing inter-species data bias towards highly investigated species.** To address the inter-species bias towards major model organisms and highly investigated species we proceed as follows. We consider the number of publications in the NCBI Pubmed database (https://www.ncbi.nlm.nih.gov/pubmed) as a proxy measure that allows us to systematically and objectively determine how highly studied (*i.e.*, "popular") a particular species is in biotechnological areas of research. In particular, we use the NCBI Entrez Programming Utilities (http://www.ncbi.nlm.nih.gov/books/NBK25501, February 2018) to obtain publication information for each of 1,840 species (Figure S7). This analysis reveals a substantial publication bias towards major model organisms and other highly studied species, indicating that current protein-protein interaction data might be prone to notable selection and investigative biases. To prevent interactomes in the long tail of less studied species to bias the main results of this study we perform all subsequent analyses on 171 species with at least 1,000 publications in the NCBI Pubmed (Figure S7 and Table S4). Furthermore, we investigate popularity of species as a possible confounding factor for interactome resilience (see Table S1 and Section S8).

We address the intra-species biases by studying various factors that could possibly confound the main results of this study and show that the relationships between evolution and interactome resilience cannot be explained by any of these biological and non-biological factors. We describe these analysis in Section S8.

#### S2 The tree of life dataset

So far, we described the interactomes used in this study. We proceed with an overview of the tree of life and phylogenetic analyses. We first discuss the phylogenetic tree of species represented in our dataset. We then describe how we extract phylogenetic taxonomy and lineage information for each species.

#### S2.1 Phylogenetic tree of species

We consider a high-resolution phylogenetic tree that we obtain based on the Hug *et al.* (*13*) study. Hug *et al.* (*13*) have constructed a phylogenetic tree that has dramatically expanded previous efforts by making use of genomes from public databases as well as newly reconstructed genomes recovered from a variety of environments. The tree includes bacteria, archaea, and eukarya and captures the diversity within each major lineage (*14*).

We here briefly overview the approach Hug *et al.* (*13*) used to construct the tree and refer the reader to (*13*) for a detailed description of the full approach. First, an alignment was generated from all SSU rRNA genes available from the genomes of the species included in the dataset. All SSU rRNA genes longer than 600 bp were aligned using the SINA alignment algorithm (*15, 16*). The full alignment was stripped of columns containing 95% or more gaps, generating a final alignment containing 1,871 taxa and 1,947 alignment positions. A maximum likelihood tree was then inferred as described in (*13*), with the RAxML run using the GTRCAT model of evolution. In particular, the RAxML inference included the calculation of 300 bootstrap iterations (extended majority rules-based bootstrapping criterion), with 100 randomly sampled to determine support values.

Hug *et al.* (*13*) note that the tree calculated using SSU rRNA gene sequence information recapitulates expected organism groupings at most taxonomic levels and is largely congruent with the tree calculated using ribosomal protein sequences. We thus use this phylogenetic tree for all the analyses in this study. In particular, we use the phylogenetic tree to characterize *evolution* for each species. Given a species $s$, its evolution $t_s$ is calculated as the total branch length (*i.e.*, nucleotide substitutions per site) from the root of the tree to the leaf representing species $s$. We establish correspondences between leaf taxa in the tree and species, for which we have interactomes (Section S1), using the NCBI Taxonomy database, which we describe next.

We share the processed data with the community with this publication (http://snap.stanford.edu/tree-of-life).

##### S2.2 The NCBI Taxonomy database

Phylogenetic taxonomy information, species names, and taxonomic lineages for all species in our dataset are extracted from the NCBI Taxonomy database (https://www.ncbi.nlm.nih.gov/taxonomy) (*17*). The taxonomy database is manually curated by a group of scientists at the NCBI who use the current taxonomic literature to maintain a phylogenetic taxonomy for organisms represented in the sequence databases. The data was accessed programmatically through the NCBI Taxonomy Browser and was processed in August 2016. Species were identified by their Taxonomy IDs. For example, *H. sapiens*, *S. cerevisiae*, *M. musculus*, and *D. discoideum* are assigned Taxonomy IDs 9606, 4932, 10090, and 44689, respectively (See Table S4 for more details). The obtained information[2] for each species include the species' common name and synonyms, full lineage information, published genome sequence information (*i.e.*, sequences represented in the NCBI nucleotide and protein sequence databases), the list of all domains within the NCBI Entrez system, and the list of various external information resources that are species-specific (*i.e.*, the NCBI LinkOut record).

We share the processed data with the community with this publication (http://snap.stanford.edu/tree-of-life).

#### S3 Information on clusters of orthologous genes and protein families

Information on protein families is extracted from an updated and extended version of the COG (Clusters of Orthologous Groups (*18, 19*)) database, which is maintained at the eggNOG (Evolutionary Genealogy of Henes: Non-supervised Orthologous Groups) (*20*). The eggNOG extends the COG methodology (*21*) to produce genome-wide orthology inferences, which are further adjusted to provide lineage-specific resolution. In particular, the eggNOG relies on UniProt/Swiss-Prot (*22*), Ensembl (*23*), and other public databases for information on protein sequences. All obtained genomes and proteomes are subjected to quality controls that prevent the inclusion of partial or draft genomes. In eggNOG, for any genomes not yet present in the COG database, or-

[2] For example, https://www.ncbi.nlm.nih.gov/Taxonomy/Browser/wwwtax.cgi?mode=Info&id=10090 (accessed in August 2018) shows the resource at the NCBI Taxonomy database for *M. musculus*.

thology assignments are made by an automatic method resembling the COG procedure. This results not only in the addition of new genes to COGs but also in the creation of a number of additional orthologous groups (*i.e.*, NOGs, non-supervised orthologous groups). Essentially, the orthology assignment procedure is based on an all-against-all pairwise Smith-Waterman comparison (*24*) of protein sequences from the selected species. The comparison uses these Smith-Waterman alignments and compositional adjustment of the scores, as in BLAST, to prevent spurious hits between low-complexity sequence regions. It also allows for recent duplications within the genome and includes a clean-up step to join remaining genes by simple bidirectional hits. Using this information on shared protein sequences, proteins are then categorized into protein families.

We use the eggNOG to compile a dataset of protein families involving species considered in this study (Section S1). A *protein family* is defined as a set of orthologous proteins (protein-coding genes) spanning multiple species. In total, we obtain 2,224 protein families, with an average of 38 proteins originating from 12 species in each family. Altogether, 81,673 distinct proteins are involved in these families.

We share the processed data with the community with this publication (http://snap.stanford.edu/tree-of-life).

#### S4 Information on natural environments of species

To study the relationship between interactome resilience and ecology of species we compile a dataset of ecological characteristics for a large number of bacterial species. The dataset covers 287 species out of 1,539 bacterial species with some available interactome data (Section S1).

We use a previously described dataset about ecology for 287 species (*25*). Freilich *et al.* (*25*) used a combination of the reverse ecology framework to examine ecological strategies for coping with competition across the microbial tree of life. First, Freilich *et al.* (*25*) calculated the biochemical environments of species using the seed set framework. Next, they simulated the expected biosynthetic capacity of each species in every such environment using the network expansion framework. Species was considered viable in a given environment if a set of essential metabolites were producible and found in the scope of the expanded network. From these data, two measures were calculated for each species, which we use in our study:

(1) *Co-habitation index:* The co-habitation (CHS) index of a species denotes the number of other species that co-populate each viable environment of the given species. This index serves as an indication of the level of competition encountered by a species in its habitats. The index focuses on each species' most populated niche (*i.e.*, maximal-CHS) representing the maximal level of competition a species encounters.

(2) *Environmental scope index:* The environmental scope index of a species is defined as the fraction of environments in which the species is viable. The index approximates environmental flexibility of a species; species with high scores are generalists that can survive in a wide span of environments, whereas species with low scores are specialists.

For each species, we also consider the following three environmental characteristics:

(1) *The fraction of regulatory genes:* The fraction of regulatory genes is the fraction of transcription factors out of the total number of genes in the organism. This is an established indicator of environmental variability of species' habitats (*26*).

(2) *Oxygen requirement:* Bacterial species are divided into three groups (aerobic, anaerobic, and facultative) according to their oxygen requirements. Oxygen-dependence annotations are retrieved from (*25*) and the NCBI's Entrez Genome Project database (http://www.ncbi.nlm.nih.gov/genomes/lproks.cgi).

(3) *Ecological habitat:* Bacterial species are divided into five groups according to their natural environments. Natural environments are categorized and ranked in the decreasing order of the environmental complexity: terrestrial, multiple, aquatic, specialized, and host-cell habitats (*i.e.*, host-associated) (*27*). Annotations for environmental complexity are retrieved from (*25*) and the NCBI classification for bacterial lifestyle (http://www.ncbi.nlm.nih.gov/genomes/lproks.cgi).

We share the processed data with the community with this publication (http://snap.stanford.edu/tree-of-life).

### S5 Additional information on interactome resilience

Next, we describe in detail our methodology for calculating the resilience of interactomes.

#### S5.1 Motivation and overview of the approach

**Motivation for the approach.** Our objective is to evaluate the topological stability, *i.e.*, robustness, of interactomes to network failures. Central to this objective is to improve our understanding of the effects of the failure of individual interactome components on the performance of the whole interactome. Here, we focus on how the *network structure* of an interactome changes as it is degraded through the removal of proteins/nodes. The motivation for studying these effects is a fundamental observation that when a network is so fragmented by the removal of nodes that the largest connected part of the network is sufficiently small (*e.g.*, only 10% of the size of the original network), then any sensible dynamical process will be unable to function on the fragmented network in an effective way (*28–30*). For example, the removal of even a small number of proteins can completely fragment the interactome and lead to cell death and disease (*31–33*) (see Section S5.4 and Table S3). The precise degree to which an interactome continues to function as individual proteins which constitute it are degraded typically depends on key features of the dynamics of the interactome. To reveal these key features it is crucial to understand the topological stability of the interactome and its resilience to failure of protein-protein inteactions. For example, the resilience of the interactome to network failures might reveal how the organism can continue to function when faced with mutations, environmental change, and internal noise, and how the organism can acquire novel properties during evolution (*34*).

**Overview of the approach.** To quantify the resilience of a given interactome and to compare the resilience of many interactomes with different sizes and connectivities we need an approach that addresses the following two challenges:

(*a*) First, the approach needs to be sensitive to subtle changes in the network structure in the sense that it can capture situations in which the network suffers a significant damage without completely collapsing.

(*b*) Second, the approach needs to take into account the size and connectivity of the original network so that resilience values of networks with different sizes can be compared.

Methodologically, our approach uses network science to quantify resilience of interactomes for all species in our dataset. We describe the interactome of each species by the connectivity of its connected components, *i.e.*, subnetworks in which any two nodes/proteins can reach each other by a path of edges/interactions. When a certain fraction of proteins out of all proteins in the species' interactome fail and are removed from the interactome we measure how the interactome becomes fragmented and how this fragmentation affects the interactome connectivity. We characterize the fragmentation by modifying Shannon diversity, which is a well-established and popular diversity index in the literature (*35–38*). We vary network failure rate and for each given failure rate analyze the fragmented interactome. The final resilience value then represents the topological stability of the interactome across all possible failure rates. We use this approach to obtain the resilience of every interactome in the protein-protein interaction dataset.

In the following subsections, we describe in detail our methodology.

**Related work on robustness of complex networked systems.** The study of the effects of the failure of individual components on the performance of a whole networked system has received considerable attention in recent years (*e.g.*, (*28–30, 39*)). The detailed motivation for studying these effects depends on the particular networked system under consideration. For example, it is clearly important to know how the failure of routers on the Internet affects the overall function of the network (*28*). Similarly, if the network in question is a contact network on which a disease can spread, then it is critical to understand how the removal of nodes from the network (*e.g.*, through vaccination) affects the spread of the disease (*40*). A common measure for robustness of networks is the percolation threshold (transition), which is defined in terms of the critical fraction of failures at which the systems completely collapses (*28*). However, Schneider *et al.* (*30*) showed that this measure may not be useful in many realistic cases. This measure, for example, ignores situations in which the network is significantly fragmented but still keeps its integrity. Besides the percolation threshold, there are other robustness measures based, for example, on the shortest path (*29*), on the graph spectrum (*41*) or on the size of the largest connected component (*30*). These measures are, however, less frequently used for being too complex or less intuitive (*30*). Furthermore, they ignore situations in which the network suffers a big damage without becoming completely fragmented and are unable to measure network fragmentation across all possible failure rates.

##### S5.2 Modified Shannon diversity

We first describe how we measure fragmentation of a given interactome at a particular network failure rate. For this we use a well-established Shannon diversity index (*35–38*), which is also known as the Shannon-Wiener index, the Shannon-Weaver index, or the Shannon entropy, which we modify to ensure that the resilience of interactomes with different numbers of proteins can be compared. In the next section, we describe how we integrate these measurements across all possible failure rates and obtain the final value for interactome resilience.

Let us consider the interactome/network $G^{(s)} = (V^{(s)}, E^{(s)})$ of species $s$ with $N$ proteins/nodes $V^{(s)}$ and $M$ interactions/edges $E^{(s)}$. Here, species $s$ is any given species from our dataset (Section S1). Let $f \in [0, 1]$ denote *network failure rate*. This rate represents a fraction $f$ nodes out of the total number of nodes in the network whose all interactions undergo a failure. That is, $f = 0$ represents a situation when all of the nodes function properly and there are not any failures, and, conversely, $f = 1$ represents a situation when all nodes fail and the network becomes completely fragmented. Upon failure of a particular node all of its interactions disappear, they are removed from the network, and the node is isolated from the rest of the network. Determining which nodes exactly will fail depends on a particular node removal strategy. This study studies resilience of interactomes under the removal of nodes uniformly at random (*i.e.*, a strategy representing random mutations in the context of biology) or in decreasing order determined based on some external information source (*i.e.*, a gene list containing information about gene essentiality). See Figure S1 for a detailed example.

When network $G^{(s)}$ is subjected to a failure rate $f$ it gets fragmented into a number of isolated network components of varying sizes (Figure S1). We quantify the connectivity of the fragmented network $G_f^{(s)}$ by calculating Shannon diversity (*35, 37*) on the resulting set of isolated components. In particular, let $\{\mathcal{C}_1, \mathcal{C}_2, \ldots, \mathcal{C}_k\}$ be $k$ isolated components in the fragmented network $G_f^{(s)}$. Let $C_i$ be the size of component $\mathcal{C}_i$, $C_i = |\mathcal{C}_i|$, *i.e.*, $C_i$ is the number of nodes that belong to $\mathcal{C}_i$. We first calculate the entropy of the resulting set of components:

$$H(G_f^{(s)}) = -\sum_{i=1}^{k} p_i \log p_i, \tag{1}$$

where $p_i = C_i/N$ is the proportion of nodes belonging to component $\mathcal{C}_i$. Necessarily, $0 \leq p_i \leq 1$ and $\sum_{i=1}^{k} p_i = 1$. We can interpret $p_i$ as the probability of seeing a node from component $\mathcal{C}_i$ when sampling one node from the fragmented network $G_f^{(s)}$. That is, Equation 1 quantifies the uncertainty in predicting the component identity of an individual node that is taken at random from the interactome and is also known as (unnormalized) Shannon diversity (*42, 43*). Finally, to correct for differences in network sizes, we modify Shannon diversity as follows:

$$H_{\text{msh}}(G_f^{(s)}) = H(G_f^{(s)})/\log N. \tag{2}$$

The normalization factor $1/\log N$ ensures that the resilience of networks with different numbers of nodes can be compared (see Section S5.3). Equation 2 represents the final formula used in this study to characterize how interactome $G^{(s)}$ fragments at a given failure rate $f$. We refer interested readers to (*36, 44*) for a detailed discussion of entropy and diversity indices. The range of possible $H_{\text{msh}}$ values is between 0 to 1, where these limits correspond, respectively, to a connected network in which any two nodes are connected by a path of edges and a completely fragmented network in which each node forms its own component. See Figure S2 for a detailed example.

##### S5.3 Interactome resilience

So far, we described how to characterize the response of a given interactome at a particular network failure rate $f$. Next, we discuss how to measure the response of that interactome across all possible failure rates and, finally, how to calculate the resilience of species $s$'s interactome.

Given a species $s$, we determine the resilience of that species' interactome $G^{(s)}$ as follows. We vary network failure rate $f$ with a one-percent step in the whole range of possible values and for each value of $f$ evaluate modified Shannon diversity $H_{\text{msh}}$ of the fragmented network $G_f^{(s)}$ using Equation 2. That is, we calculate $H_{\text{msh}}$ as a function of failure rate $f$, which allows us to quantify how fragmentation of the network depends on the fraction of nodes removed. In particular, we start with the full network $G^{(s)}$ and $f = 0$. For each next possible value of $f = qN/N \cdot 100\%$, for $q = 0.01, 0.02, \ldots, 1$ ($N$ is the total number of nodes in the interactome of species $s$), we remove an additional one-percent of the total number of nodes uniformly at random from the current network. We then use $H_{\text{msh}}$ (Equation 2) to calculate the fragmentation of the resulting network $G_f^{(s)}$. The whole procedure is then repeated for the next value of $f$ with the resulting network as the input. The final result of this procedure is a resilience curve that represents fragmentation of the network at each possible failure rate. Because modified Shannon diversity $H_{\text{msh}}$ is normalized, the resilience curve is monotonically increasing (*i.e.*, when increasing the failure rate, the interactome can only become more fragmented), it reaches its minimum value of 0 at $f = 0$ (*i.e.*, the interactome is connected) and its maximum value of 1 at $f = 1$ (*i.e.*, the interactome is completely fragmented). See Figure S3 for a detailed example and interpretation. Finally, the *interactome resilience* for species $s$ is obtained as one minus the area under the resilience curve (Resilience $= 1 -$ AUC). Formally, the interactome resilience for species $s$ is calculated as:

$$\text{Resilience}(G^{(s)}) = 1 - \int_0^1 H_{\text{msh}}(G_f^{(s)})\, \mathrm{d}f. \quad (3)$$

The interactome resilience thus takes values between 0 and 1; a higher value implies a more resilient interactome.

We just described our approach (Section S5.2-Section S5.3) for calculating the interactome resilience of a particular species $s$. We note that we use this methodology to calculate the resilience of every interactome in our dataset. The interactome resilience values for all species are shown in Table S5.

##### S5.4 Removal of nodes representing essential protein-coding genes

Network failures as described above represent a situation in which randomly selected proteins from the interactome fail (*e.g.*, by random mutations or environmental factors such as availability of resources). Apart from eliminating the proteins randomly, another particularly interesting procedure is to remove proteins in the order determined based on essentiality information. Such a procedure represents an adversarial agent that attempts to deliberately damage the interactome by preferentially targeting proteins that have a vital role in the survival of the organism (*32*). To investigate how vulnerable the interactomes are to these targeted attacks (*28*) we conduct a series of additional analyses. First, we identify six species in our dataset (*i.e.*, humans, *S. cerevisiae*, *M. musculus*, *D. melanogaster*, *C. elegans*, *A. thaliana*) for which we obtain genome-wide information on gene essentiality (*i.e.*, whether a particular protein-coding gene is essential or not). We then use our methodology (Section S5.2-Section S5.3) together with this information to calculate a resilience value for each interactome. As proteins are selected and removed from the interactome based on whether they are encoded by essential genes, the calculated resilience values represent the attack vulnerability of the interactomes. That is, a lower value indicates a greater vulnerability of an interactome to attacks on essential genes.

**Results.** Results of these analyses are shown in Table S3. Across six species, including humans, *S. cerevisiae*, *M. musculus* and others for which genome-wide essentiality information exists, we find that interactomes are significantly less resilient to failures of essential genes than to failures of random genes ($p$ value $< 1 \cdot 10^{-4}$; permutation test). This finding demonstrates that interactomes have a topological structure that is error-tolerant but extremely vulnerable to targeted attacks on essential genes. That is, when essential genes are targeted, a typical interactome becomes rapidly fragmented and breaks into many small isolated components. This decrease in resilience provides evidence for the topological instability of interactomes to targeted attacks on essential genes. We note that these results are in agreement with our current understanding that essential genes tend to encode for proteins that play a vital role in maintaining the interactome' connectivity (*32, 45*). These results also provide further empirical motivation for our study of interactome resilience as failures of proteins can affect the interactome to the extent that the interactome loses its biological function and the disrupted interactions increase the risk of diseases (*33*).

#### S6 Additional information on analysis of protein network neighborhoods

Next, we present a detailed discussion of our methodology for the analysis of protein network neighborhoods and describe the different statistical tests and controls.

##### S6.1 Protein network neighborhoods

For each species, we construct a separate protein network neighborhood for every protein in that species' interactome. Consider the interactome $G^{(s)}$ of species $s$ and a protein/node $u \in V(G^{(s)})$ that is part of the interactome. The $u$'s network neighborhood $N_k(u)$ is a *centered graph* (*46*). In particular, $N_k(u)$ is centered at $u$ and is a subgraph of the interactome $G^{(s)}$ that consists of $u$, its $k$-hop neighbors in the interactome, and all of the interactions/edges between them. See Figure S4 for a detailed example.

Protein network neighborhoods are thus built around a particular protein designated as a central node $u$. To construct a network neighborhood for a particular protein in a particular species, we begin by taking all the other proteins with whom the central protein $u$ interacts, either directly (if $k = 1$) or both directly and indirectly (if $k > 1$) Finally, we note all interactions among those other proteins. The result is a mini-network or $k$-hop neighborhood surrounding $u$ that can reveal something biologically meaningful from $u$'s perspective. Motivated by previous observations (*47, 48*) that first- and second-order neighbors are most informative of individual proteins, we use in this study $k = 1$ and $k = 2$. There are several more or less obvious but interesting properties of protein network neighborhoods, which follow from the centered graph theory:

(1) In terms of density, protein network neighborhoods have two extremals: the minimal star graph and the maximal complete graph (where all possible edges are present).

(2) Protein network neighborhoods are, of course, connected; that is, there is a sequence of nodes and edges—a path—from any node to all others.

(3) The longest shortest path linking any pair of nodes is less than or equal to $k+1$: this is called the diameter of the network neighborhood.

(4) Any shortest path connecting a pair of nodes has, according to (3) above, length equal to either $1, 2, \ldots, k+1$. Specifically, in the case of $N_1(u)$, if a path has a length of 2, the end nodes are in different components that are linked by a mid-point, which is the central node $u$. Since path lengths form a partition in a network neighborhood, we can examine the structure of these components and the pattern of connectivity by counting either one. In this study, the structure of these components, as well as the connectivity pattern are used, which we describe next.

#### S6.2 Analyses of protein network neighborhoods

We characterize each protein's network neighborhood (Figure S4) by calculating two network metrics, which we describe next. Results of these analyses are shown in Figure 3 in the main text.

**Isolated components of a protein network neighborhood.** The metric is defined as the number of connected components that arise when the central node $u$ is removed from the neighborhood and it is normalized by $u$'s degree:

$$\mathrm{IC}(u) = \frac{n}{d_u}, \tag{4}$$

where $n = |\{\mathcal{C}_1, \mathcal{C}_2, \ldots, \mathcal{C}_n\}|$ represents the number of isolated network components in the fragmented version of network neighborhood $N_2(u)$ and $d_u = |N_1(u)|$ represents node $u$'s degree. Note that $n$ is always bounded from above by $d_u$. The metric thus takes values in $[1/\max_u d_u, 1]$ and a higher value indicates a greater fragmentation of the protein network neighborhood. See Figure S5 for a detailed example.

**Effective size of a protein network neighborhood.** We start with a brief overview of the concept of structural holes in network science and then proceed with the definition of the network metric. *Network structural holes* are "gaps" that exist between different areas of a network, that is, network areas that have few edges/interactions between them. The foundational work of this concept (*49*) highlights network structural holes as a mechanism that, at a local level, can be seen as a separation between nonredundant nodes within a given network neighborhood (Figure S5). In particular, to identify network structural holes, one begins at the local level with a network neighborhood. Taking a central node $u$ of the neighborhood, a redundant neighbor of $u$ is one that is also connected to other neighbors of $u$. This means that when $u$ is connected to non-redundant neighbors, $u$ sits on a "bridge" between those separate areas of its local network neighborhood. The task of identifying structural holes is thus a matter of identifying neighbors of $u$ that are not connected to each other (*49–51*).

Given a central node $u$, the notion of redundancy captures the extent to which another node $v$ in $u$'s neighborhood is related to some third node $w$ that is also a part of $u$'s neighborhood. Such neighbors are redundant to the extent that they lead to the same nodes, and so provide similar functional/information benefits (*49*). Gaining a handle on which neighbors of $u$ are redundant in $u$'s neighborhood helps us understand the extent to which $u$ is connected to disparate or unconnected (*i.e.*, non-redundant) neighbors, and thus it illuminates the bridging potential of $u$ in the network. This bridging potential is captured by the *effective size* of $u$'s neighborhood or the "true" size of $u$'s network absent of redundant neighbors.

Following (*49*), we define effective size mathematically as follows. The effective size of $u$'s network neighborhood is the sum of the non-redundant portion of $u$'s connections over all $u$'s neighbors:

$$\mathrm{ES}(u) = \sum_{v \in N_1(u)} (1 - 1/d_u \sum_{w \in N_1(u)} e_{vw}) = d_u - 1/d_u \sum_{v \in N_1(u)} \sum_{w \in N_1(u)} e_{vw}), \quad w \neq u, v \tag{5}$$

where $e_{vw} = 1$ if nodes $v$ and $w$ are connected and is 0 otherwise. Note that the first summation covers all neighbors $v$ in $u$'s local network, and the second sum covers all intermediary connections $w$ between $u$ and $v$. Note also that ES is always bounded from above by $d_u$ and that ES achieves the maximum value exactly when $u$'s network neighborhood $N_1(u)$ is a star graph. We thus normalize the metric by dividing its value by $d_u$, so that the metric takes values in $[0, 1]$, and that a higher value indicates a larger effective size of the network neighborhood. See Figure S5 for a detailed example.

### S7 Additional information on analysis of interactome networks

Next, we present a detailed discussion of our network-based methodology and describe the different statistical tests and controls.

#### S7.1 Protein-protein interaction rewiring rates (IRR)

We develop an approach to quantify protein-protein interaction rewiring based on analogy to simple models of sequence evolution and use it to conduct a systematic study on all the interactomes. Results of these analyses are shown in Figure 4 in the main text. Next, we describe the approach.

**Calculating interaction rewiring rates (IRR).** We use a consistent method to calculate interaction rewiring rates comparing protein network neighborhoods of two orthologous proteins across species. First, orthology relationships between proteins/nodes in species are established (Section S3). For a network motif $m$ of interest (*e.g.*, a simple edge/interaction, a triangle, a square), we then count the number of instances of that network motif in each of the two compared protein network neighborhoods. We use these counts to then calculate the fraction of possible instances of $m$ that exist in the neighborhood. For example, when $m$ is a triangle involving the central node $u$, this gives us the probability that two neighbors of node $u$ are connected with each other (*i.e.*, triangle clustering (*52*)). In another example, when $m$ is a square touching the central node $u$, this gives us the probability that two neighbors of node $u$ share a common neighbor different from $u$ (*i.e.*, square clustering (*53*)). This means that these calculated values are directly comparable between protein network neighborhoods, even if neighborhoods are of different sizes and connectivities. Finally, the following equation is used to calculate the *interaction rewiring rate* for a pair of protein network neighborhoods:

$$\text{IRR}(m; u, v) = \log_2 \frac{m(u)}{m(v)}, \tag{6}$$

where $u$ and $v$ are the proteins whose neighborhoods are compared. Here, $m(u)$ ($m(v)$) denotes the fraction of possible instances of $m$ that exist in the neighborhood of $u$ ($v$), that is, it represents the probability of observing $m$ in the neighborhood of $u$. Here, given two proteins from different organisms, $u$ is selected as the protein from the organism with more nucleotide substitutions per site, *i.e.*, $t_{\text{species}(u)} > t_{\text{species}(v)}$. Interaction rewiring rate $\text{IRR}(m; u, v)$ thus measures the fold change between the probability of $m$ occurring in the neighborhood of protein $u$ relative to the probability of the same motif occurring in the neighborhood of an evolutionarily younger orthologous protein $v$. A IRR value greater than 0 means that the motif $m$ becomes more abundant with the evolution and vice versa. We compute the rates IRR for all orthologous protein pairs and summarize the values by reporting the mean, median, and other statistics.

Importantly, Equation 6 specifies an *instantaneous* rewiring rate, which we use to compare networks between *closely related species* (*i.e.*, $t_{\text{species}(u)} - t_{\text{species}(v)} < 0.1$). We note that the instantaneous rewiring rate is preferred over average rewiring rate (*i.e.*, $\text{IRR}(m; u, v) = \log_2(m(u)/m(v))/(t_{\text{species}(u)} - t_{\text{species}(v)})$) because of the following reasons (*54*). For evolutionarily distant species, network rewiring approaches saturation and is hard to compare. This is because new network structural changes happen on top of previous changes, which then have only little effect on the rewiring. In particular, Shou *et al.* (*54*) found that similar to nucleotide sequences (*i.e.*, Jukes-Cantor model), biological networks show a decreased rate of change at large evolutionary distances because of saturation in potential substitutions. For these reasons, the interaction rewiring rates in this study are based on the comparison of networks between *closely related species* using instantaneous rewiring rates.

**Conducting randomization-test procedures.** To evaluate the statistical significance of the obtained values of IRR we use two complementary random models:

(1) *Randomized evolutionary distances:* For network neighborhoods of two orthologous proteins $(u, v)$, randomize information on the evolution of each protein's originating species. This means that the protein $u$ in the numerator of Equation 6 can sometimes come from the organism with fewer nucleotide substitutions per site than the protein $v$ in the denominator of Equation 6, *i.e.*, $t_{\text{species}(u)} < t_{\text{species}(v)}$. This model tests for whether interactions rewire in a way that is independent of species' evolution, *i.e.*, the amount of genetic change a species has undergone.

(2) *Randomized orthologous relationships:* First, randomize orthologous relationships between proteins. Use the new randomized relationships to calculate interaction rewiring rates as described in the previous paragraph. This model tests for whether there exists a global mechanism, which is independent of orthologous relationships, that determines how interactions rewire.

The statistical significance of an observed difference between the values of IRR and the randomized counterpart $\text{IRR}_{\text{randomized}}$ is given by the $p$ value from a two-sample Kolmogorov-Smirnov test.

#### S7.2 Interactome network null models

We also explore whether the resilience of interactomes could be the result of a particular structure intrinsic to interactome networks (*e.g.*, (*1*)). Using the configuration model (*55, 56*) we construct 171,000 randomized versions of the interactomes, *i.e.*, 1,000 randomized interactomes for each of 171 species. The randomized interactomes have the same number of proteins/nodes and interactions/edges, and the same node degrees as the actual interactomes, but have randomized interactions (*i.e.*, degree preserving randomization (*57*)).

**Results.** We find that 171 out of 171 species have interactomes whose resilience is statistically significant with respect to the random expectation, hence the interactome resilience cannot be attributed to structural network properties alone. We also compare the distribution resilience for bacterial and eukaryotic interactomes with the distribution of resilience for random interactome counterparts. Again, we find that the observed differences between the interactomes and their randomized counterparts cannot be explained solely by network size and degree distribution (Figure S9).

##### S7.3 Estimating the size of the whole human interactome

Next, we describe how our network-based methodology can be used to estimate the size of the whole human interactome from the currently available (incomplete) network data. The actual size of the whole human interactome is currently unclear and its estimation is a highly non-trivial task. It will likely remain so until most of the interactome becomes accessible to experimental technologies and we get a fairly complete description of the interactome. The interest in knowing the size of the whole human interactome, *i.e.*, the number of protein-protein interactions in humans, stems from a surprising result of genome-sequencing projects that the number of genes in species as diverse as fruit flies, nematodes, and humans does not reflect our perception of their relative complexity (*58*). In other words, very different organisms have a surprisingly similar number of genes (*59*). For example, *C. elegans* has a similar number of genes as humans, whereas rice and maize have even more genes than humans. It was quickly suggested that the biological complexity of organisms is not reflected merely by the number of genes but by the number of physiologically relevant interactions and that the structure of interactome is one of the crucial factors underlying the complexity of organisms (*58, 60, 61*).

To address this challenging question, many studies attempted to provide statistical estimates of the size of the whole human interactome based on currently available partial subnet data (*e.g.*, (*11, 58, 62–64*)) and show that the estimated sizes correlate much better with the apparent biological complexity of different organisms. In contrast to some previous studies (*e.g.*, (*58, 63*)), the main objective of our study was not to develop a new statistical procedure to estimate the size of the whole interactome. Nevertheless, we are able to obtain an estimate of the size of the whole human interactome that is in surprisingly good agreement with the previous estimates by simply using interaction rewiring rates (IRR, see Section S7.1).

**Interactome size as a by-product of interaction rewiring rates (IRR).** In addition to using IRR, our approach uses information about a reference organism, which can be any organism for which a fairly extensive description of the interactome is available. We use *S. cerevisiae* as a reference organism as we are beginning to have a fairly complete description of its interactome. To estimate the interactome size of a given organism, we then extrapolate interactome size of the reference by using two pieces of information about the organism of interest: the number of protein-coding genes and the evolution of the organism (as defined in Section S2). Note that the latter information is available for any organism with a completed genome sequencing project. Next, we discuss in detail the method for estimating the size of the whole interactome.

**Calculating an estimate of the human interactome size.** Our goal is to calculate an estimate for the number of protein-protein interactions in humans. We take *S. cerevisiae* as a reference organism. Let $t_y = 3.736$ indicate the evolution of yeast (Section S2), $N_y = 5{,}800$ represent the number of yeast genes (*63*), and $M_y = 52{,}500$ represent the mean projected number of interactions in the yeast interactome (see Table 2 in (*63*)).

To estimate the human interactome size, we need the number of human protein-coding genes and characterization of the evolution of humans. Let $t_h = 3.997$ indicate the evolution of humans (Section S2) and let $N_h = 20{,}400$ represent the number of human protein-coding genes based on the number of protein-coding genes in the Ensembl GRCh38.p12[3]. Next, we use these data to calculate the number of protein-protein interactions in humans, $M_h$.

Our method makes two important assumptions that can be relaxed without any change to the approach as a whole. Following Stumpf *et al.* (*58*) and taking into account present experimental methods, we ignore multiple splice variants per gene. Second, to make the method applicable to as many organisms as possible we want to use an unconstrained network model of edges/interactions. We use the Erdös-Renyi model, which is defined as a random graph with $N$ nodes where each possible edge has a probability $p$ of existing. The model does not impose any constraints on how the edges are distributed among the nodes. We make use of two well-known

[3]This genome assembly corresponds to GenBank Assembly ID GCA_000001405.27. The assembly is available at http://www.ncbi.nlm.nih.gov/genome/assembly/?term=GCA_000001405.27.

facts about Erdös-Renyi graphs (*65*): the expected number of edges in an Erdös-Renyi graph is: $M = \binom{N}{2}p$, and its expected mean degree is: $A = Np$. In what follows, we use the subscript to denote organism name.

The calculation consists of four steps. First, we take $M_y$ and $N_y$ and compute the probability $p_y$ of each possible edge/interaction in yeast as: $p_y = M_y/\binom{N_y}{2} = 0.00312$. Second, we calculate the expected mean degree in the yeast interactome as: $A_y = N_y\, p_y = 18.107$. Third, we use the expected mean degree $A_y$ and the rewiring rate of an individual edges/proteins, IRR $= -0.215$ (Section S7.1 and Figure 4), to calculate the expected mean degree $A_h$ in the human interactome as: $A_h = 2^{\mathrm{IRR}(m_1)} A_y = 15.600$. We use the resulting $A_h$ to compute the probability $p_h$ of each possible edge/interaction in human as: $p_h = A_h/N_h = 0.000765$. Finally, we obtain $M_h$ as: $M_h = \binom{N_h}{2}p_h = 159{,}109$. Thus, the projected human interactome size is approximately 160,000 interactions.

Taken together, the projected interactome size is generated by a simple, very approximate, but surprisingly effective, statistical arguments that extrapolate the yeast interactome to the human interactome. However, this prediction is in surprisingly good agreement with three previous estimates of the size of the human interactome (*62–64*), which range from $M_h = 150{,}000$ to $M_h = 370{,}000$ interactions and are generated by rather involved statistical procedures.

### S8 Additional analyses on possible confounding factors

Next, we present a detailed discussion of our statistical methodology and describe how we control for possible confounding factors when determining the relationship between evolution and interactome resilience (see Figure 1 in the main text).

#### S8.1 Confounding factors and partial correlation analyses

One key question is whether our main results could be an artifactual finding arising due to the uneven size of interactome networks, broad-tailed degree distributions, the presence of high-degree nodes (hubs), or other network structural and genomic properties of species. To answer this question, we design a causal model (Figure S8) that we use to systematically study alternative hypotheses that could potentially explain the relationship between evolution and interactome resilience.

Next, we describe in detail the statistical procedures used to perform these analyses. Evolution and interactome resilience (shown in Figure 1) are correlated, however, it is difficult to say why this relationship exists. One reason for the difficulty is the likely presence of *confounding factors*. In particular, some third variable, a confounder $Z$, may be producing changes in both evolution and interactome resilience and thus could lead to artifactual findings. In what follows, we describe the statistical procedures that allow us to show that the relationship between evolution and interactome resilience is not confounded by any of the several possible confounders listed in Figure S8.

Let $E$ represent a vector of evolutionary information (*i.e.*, nucleotide substitutions per site) for all the species in our dataset (Section S2) and let $R$ represent a vector of corresponding interactome resilience values. Additionally, let $Z$ denote a vector of values of a particular confounding factor (*e.g.*, the number of interactions/edges in each interactome; see Figure S8 and Table S1 for the full list of confounding factors). Partial correlation is a procedure that uses multiple regression to determine what the correlation between $E$ and $R$ would be (hypothetically) if they were not each correlated with the third variable, a possible confounding factor $Z$. Alternatively, we say that partial correlation allows us to determine what the correlation between $E$ and $R$ would be (hypothetically) if the third variable $Z$ were held constant. More specifically, we use the following statistical procedures:

(1) *Parametric partial measure of association:* We use the linear regression approach (*66*) to compute the partial correlation $r_{ER|Z}$. In particular, taking $E$ and $R$ and a possible confounding factor $Z$ the algorithm

can be summarized as follows: 1) perform a linear least-squares regression with $E$ as the target and $Z$ as the predictor, 2) calculate the residuals in Step #1, 3) perform a linear least-squares regression with $R$ as the target and $Z$ as the predictor, 4) calculate the residuals in Step #3, and 5) calculate the correlation coefficient between the residuals from Steps #2 and #4. The result is the partial correlation $r_{ER|Z}$ between $E$ and $R$ while controlling for the effect of $Z$.

(2) *Nonparametric partial measure of association:* We consider partial rank correlation (Spearman's partial $\rho$) (*67–69*) to compute the partial rank correlation coefficient $\rho_{ER|Z}$ between evolution $E$ and interactome resilience $R$ given the effect of a confounding factor $Z$. Partial rank correlation $\rho_{ER|Z}$ is the rank correlation between $E$ and $R$ after removing the effect of $F$ and can be computed based on standard rank correlations $\rho$ between the three variables $E$, $R$, and $Z$ as follows:

$$\rho_{ER|Z} = \frac{\rho_{ER} - \rho_{EZ}\rho_{RZ}}{\sqrt{(1-\rho_{EZ}^2)(1-\rho_{RZ}^2)}} \tag{7}$$

with $\rho_{YX}$ denoting the rank correlation between $X$ and $Y$. As with the standard rank correlation coefficient, a value $\rho_{ER|Z}$ of $+1$ indicates a perfect positive linear relationship, a value $\rho_{ER|Z}$ of $-1$ indicates a perfect negative linear relationship, and a value $\rho_{ER|Z}$ of $0$ indicates no linear relationship.

**Results.** Results of these analyses are shown in Table S1. Taken together, we find that within the limitations imposed by the incomplete interactome data, the relationship between evolution and interactome resilience systematically persists when the effects of possible confounding factors are removed. For any given confounder $Z$ in Table S1 (*e.g.*, interactome density, genome size), we find that partial correlation values ($r_{ER|Z}$ and $\rho_{ER|Z}$ for any $Z$) are substantial and significantly larger than zero. This finding indicates that a significant relationship between evolution $E$ and interactome resilience $R$ exists even if we control for $Z$, that is, if we statistically hold $Z$ constant. In other words, the confounders *only partly account for* the relationship between evolution and interactome resilience and cannot explain the observed relationship. Based on that, we conclude that our main results are not direct effects of various properties of species' genomes, such as genome size and the number of protein-coding genes. Furthermore, our main results are not direct effects of various properties of species' interactomes, such as network size, the number of interactions in each species, and the presence of hubs in the interactome networks.

##### S8.2 Comparison with unbiased datasets

We complement our analysis using only interactions from well controlled and completely unbiased high-throughput yeast-two-hybrid (Y2H) datasets (*11, 31, 62, 64, 70–74*). These data are particularly suited to addressing the effects of incompleteness systematically because all possible pairwise combinations of a given set of proteins have been tested in an unbiased fashion on the same platform. We systematically explore how our main results are affected when only these high-throughput protein-protein interaction data from human and yeast are used. We note that these additional experiments do not require any change to our interactome resilience methodology (Section S5), as the methodology can be used to compare the resilience of interactomes that can be of vastly different sizes.

**Using only yeast-two-hybrid (Y2H) protein-protein interaction data.** We compiled five distinct *S. cerevisiae* protein-protein interaction datasets (*31, 70, 71, 73, 74*) and four distinct *H. sapiens* protein-protein interaction datasets (*11, 62, 64, 72*), each dataset resulting from a high-throughput yeast-two-hybrid assay. Each dataset represents an unbiased systematic screen because all pairwise combinations between a set of proteins were interrogated (*i.e.*, all pairwise interactions within a set of proteins were tested). Since interactome data are prone to investigative biases (Section S1), we use these unbiased datasets to systematically address the effects of investigative biases.

Our aim is to study how the values of interactome resilience change when only unbiased high-throughput data are used to quantify the resilience instead of the full species' interactomes (*i.e.*, data described in Section S1). To this aim, we use our interactome resilience methodology with each of these nine additional high-throughput Y2H datasets. We then compare the results obtained on these datasets with the results obtained on the full interactome data. In particular, we systematically compare each (*S. cerevisiae*, *H. sapiens*) high-throughput Y2H dataset pair with the (*S. cerevisiae*, *H. sapiens*) full interactome dataset pair. We examine whether the values of interactome resilience are consistent across these dataset pairs. That is, we ask the following question: If *S. cerevisiae* has higher interactome resilience than *H. sapiens* on the full data, does it also have higher interactome resilience when only the high-throughput Y2H data are used?

**Results.** Results of these analyses are shown in Table S2. Based on these results, we conclude that within the limitations imposed by the current protein-protein interaction data the interactome resilience continues to exist in unbiased high-throughput data (*i.e.* in 17/20=85% dataset pairs) and that our main findings can be reproduced even in much sparser/smaller high-throughput interaction datasets from Y2H.

#### Supplementary Figures

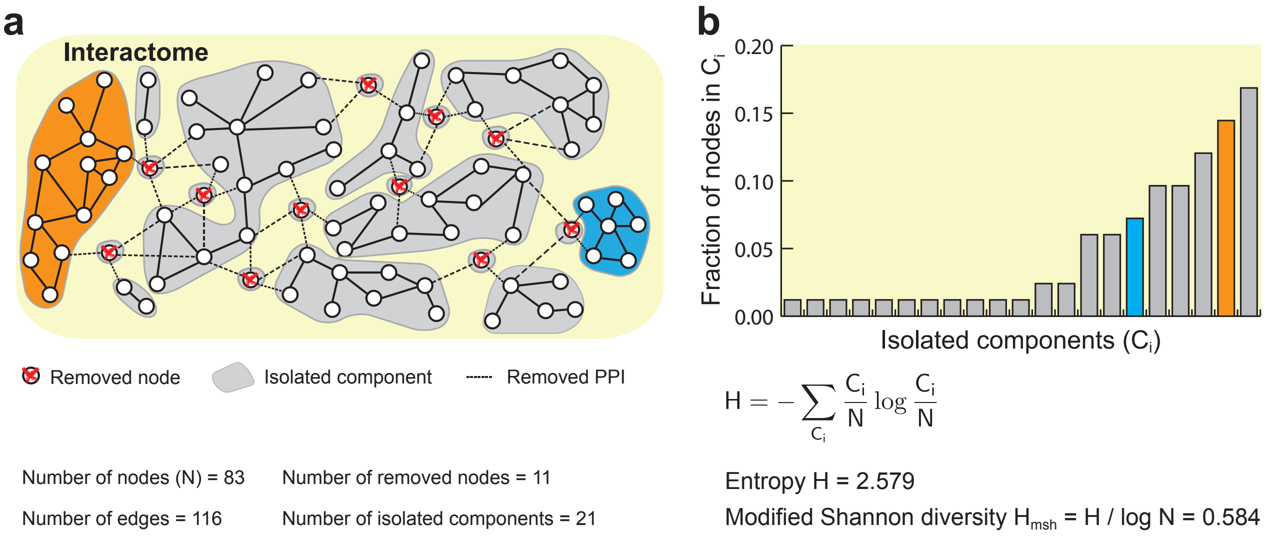


**Figure S1: Characterizing fragmentation of the interactome into isolated components upon node removal.** **(a)** Shown is a hypothetical interactome whose nodes represent proteins and edges indicate physical protein-protein interactions (PPIs). The interactome network has $N = 83$ nodes and is (initially) connected, *i.e.*, one can traverse from one node to any other node in the network following the edges. In this example, 11 nodes are selected at random and their PPIs are removed from the interactome. This results in a fragmented interactome with 21 isolated components (in grey). Highlighted are two isolated components whose sizes are $C_1 = 12$ (orange), and $C_2 = 6$ (blue). **(b)** The fragmentation of the interactome is characterized by isolated components and quantified by modified Shannon diversity $H_{\text{msh}}$ (Section S5). The plot shows the fractional size of each isolated component, *i.e.*, $C_i/N$. Using information about fractional component sizes as input to the normalized entropy formula, we obtain $H_{\text{msh}} = 0.584$.

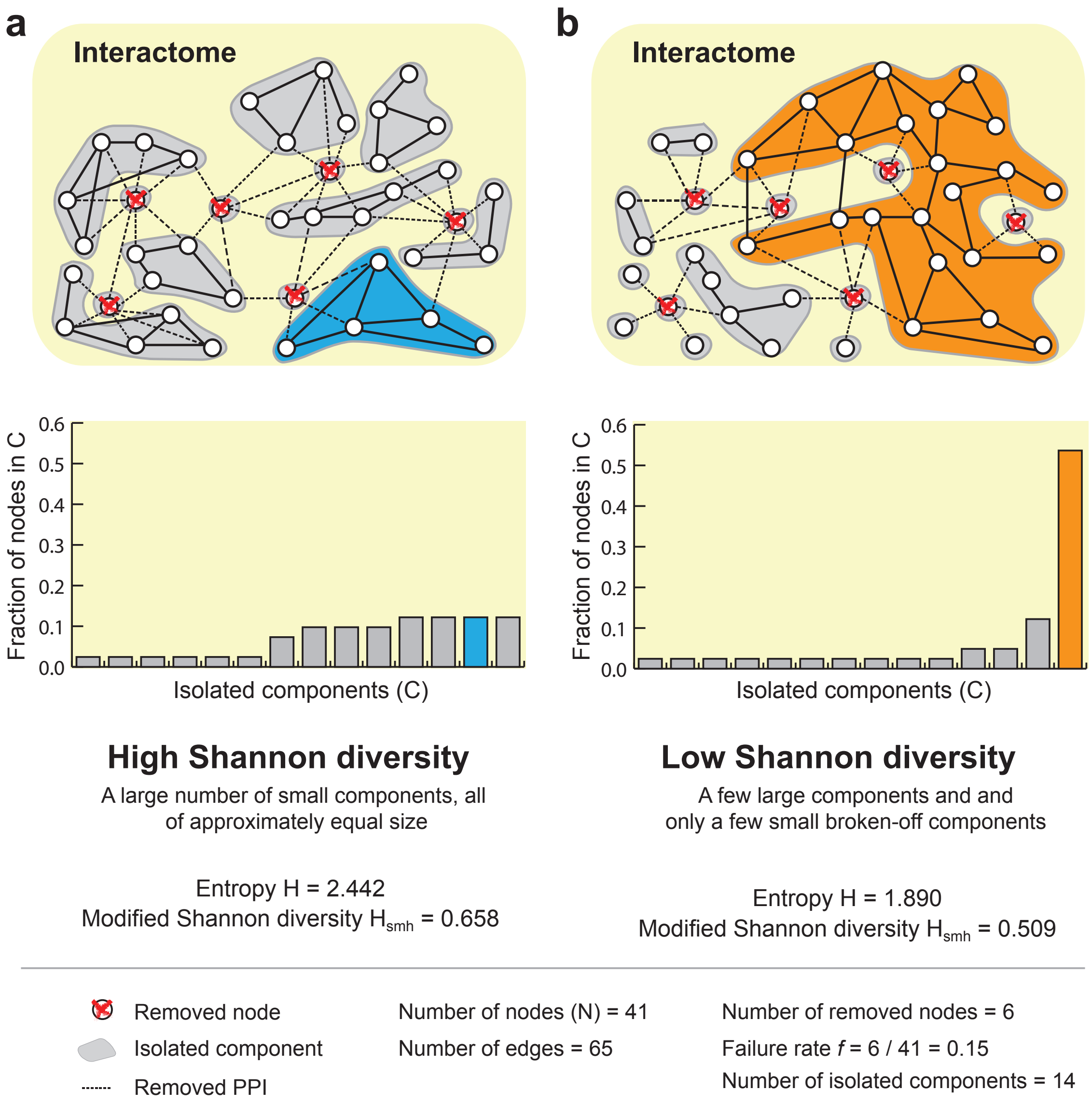


**Figure S2: Quantifying fragmentation of the interactome using modified Shannon diversity.** Graphical explanation of modified Shannon diversity, a measure used to characterize how the interactome fragments into isolated components at a given network failure rate. Shown are two hypothetical interactomes whose nodes represent proteins and edges indicate physical protein-protein interactions (PPIs). The interactome networks have the same number of nodes ($N = 41$) and the same number of edges ($E = 65$) but different connectivities, *i.e.*, edges connect different node pairs in each network. This example illustrates how interactomes with different connectivities can fragment in different ways even though they are subjected to the same failure rate (*i.e.*, the same number of nodes removed from each interactome). **(a)** In this interactome, 6 nodes are selected at random and their PPIs are removed from the interactome. The interactome get fragmented into 14 isolated components (in grey), which are relatively small and of approximately equal size. Even the largest isolated component contains less than 10% of the nodes as seen in the plot. Because of that, modified Shannon diversity $H_{\text{msh}}$ describing fragmentation of the interactome is high, $H_{\text{msh}} = 0.658$ (Section S5). **(b)** As in the previous interactome, 6 nodes are selected at random and their PPIs are removed from the interactome resulting in 14 isolated components. However, the interactome falls apart into a few large isolated components (the largest isolated component contains more than 50% of the nodes) and a few small broken-off components. Because of that, modified Shannon diversity $H_{\text{msh}}$ is lower than in (a), $H_{\text{msh}} = 0.509$.

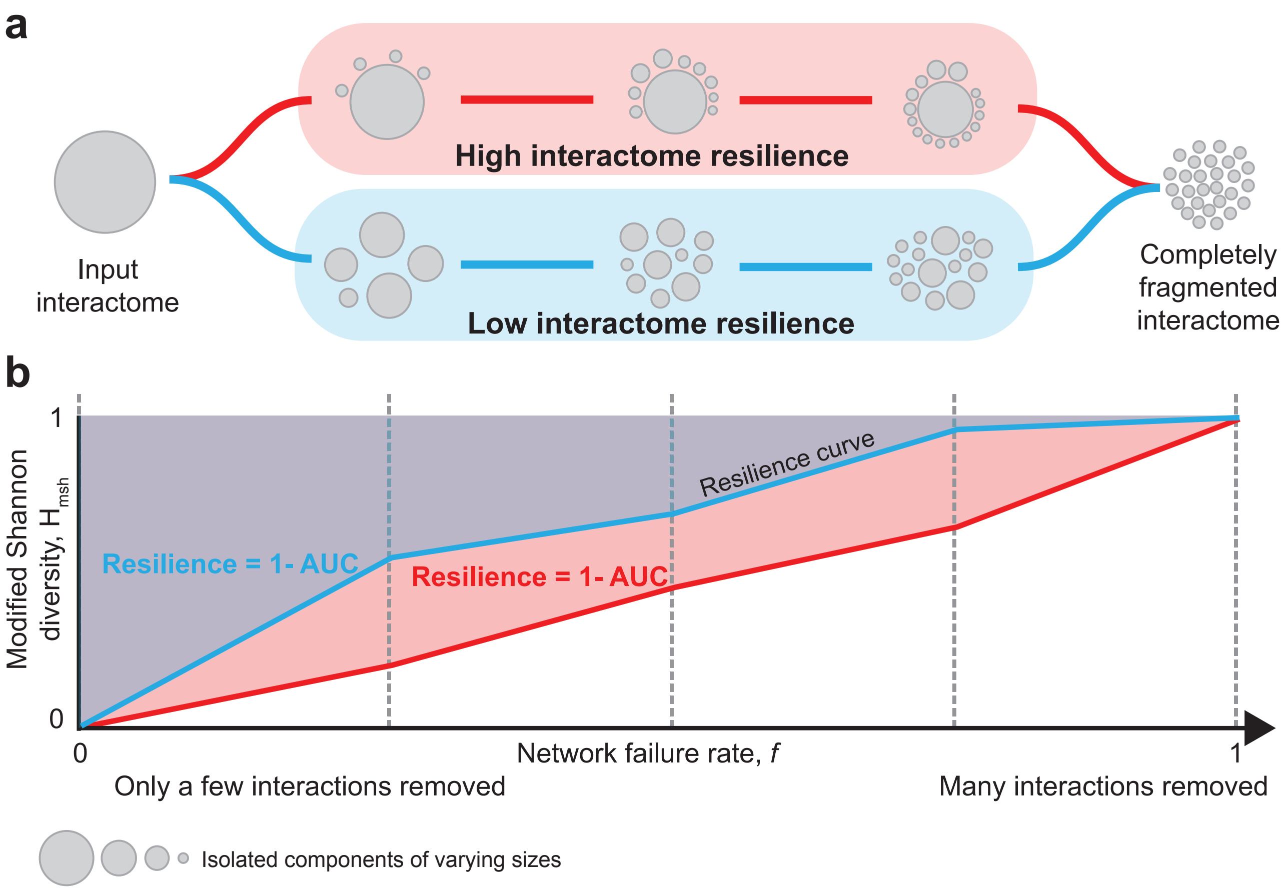


**Figure S3: Interactome resilience.** Graphical definition of interactome resilience using modified Shannon diversity (see Figure S2 for the explanation of modified Shannon diversity). **(a)** Resilience summarizes response of the interactome to failures across all possible failure rates $f$, $f \in [0, 1]$. Fragmentation of a highly resilient interactome follows the following scenario (in red; from left to right). For a small failure rate $f$, the interactome breaks into one (or, only a few) large components and a few small broken-off components. The size of the largest component slowly decreases as $f$ increases. That is, the increasing failure rate leads to the isolation of small components only and the interactome slowly deflates as these small components break off one by one. Thus, the interactome stays together as a large component for very high values of $f$, providing evidence of the topological stability of the interactome under failures. A non-resilient interactome follows a different scenario under failures (in blue; from left to right). For a small failure rate $f$, components of different sizes break off, although there are still a few relatively large components. These isolated components then quickly break into small fragments and large components completely disappear. At even higher $f$ the components are further fragmented into single nodes or components of size two. Ultimately, when $f = 1$, the interactome is completely fragmented into $N$ isolated components, each containing exactly one node. **(b)** Quantitatively, fragmentation of the interactome at each value of $f$ is calculated using modified Shannon diversity (Figure S1 and Figure S2). Repeating this calculation for various values of $f$ results in a resilience curve (Section S5). The resilience curve is monotonically increasing (*i.e.*, when increasing the failure rate, the interactome can only get more fragmented), it reaches its minimum value of 0 at $f = 0$ (*i.e.*, the interactome is connected) and its maximum value of 1 at $f = 1$ (*i.e.*, the interactome is completely fragmented). Resilience of the interactome is then obtained as one minus the area under the resilience curve (Resilience $= 1 -$ AUC). As a result, the interactome resilience takes values between 0 and 1, a higher value implies a more resilient interactome.

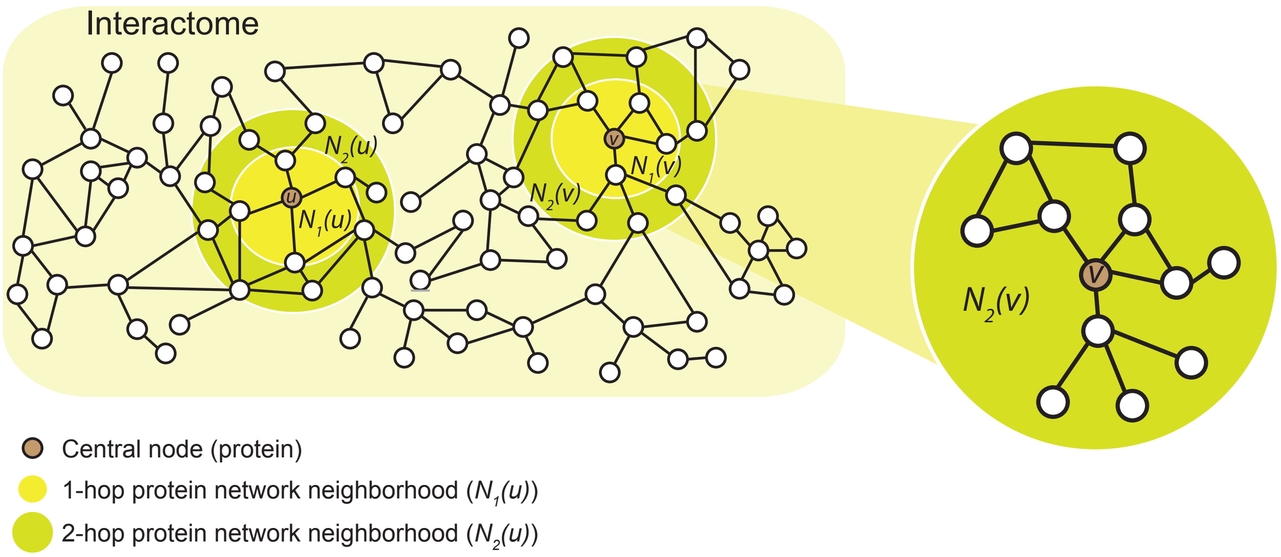


**Figure S4: Protein network neighborhoods in the interactome.** Shown is a hypothetical interactome whose nodes represent proteins and edges indicate physical protein-protein interactions (PPIs). To investigate network structural changes in local protein neighborhoods (*46*), we decompose a species' interactome into local protein networks, using a 2-hop subnetwork centered around each protein in a given species (*i.e.*, $N_2(u)$ for node $u$). The subnetwork is then used as a local representation of the protein's direct and nearby interactions in the species' interactome (see also Section S7). Highlighted are 1-hop (in yellow) and 2-hop (in green) protein network neighborhoods for two nodes, $u$ and $v$.

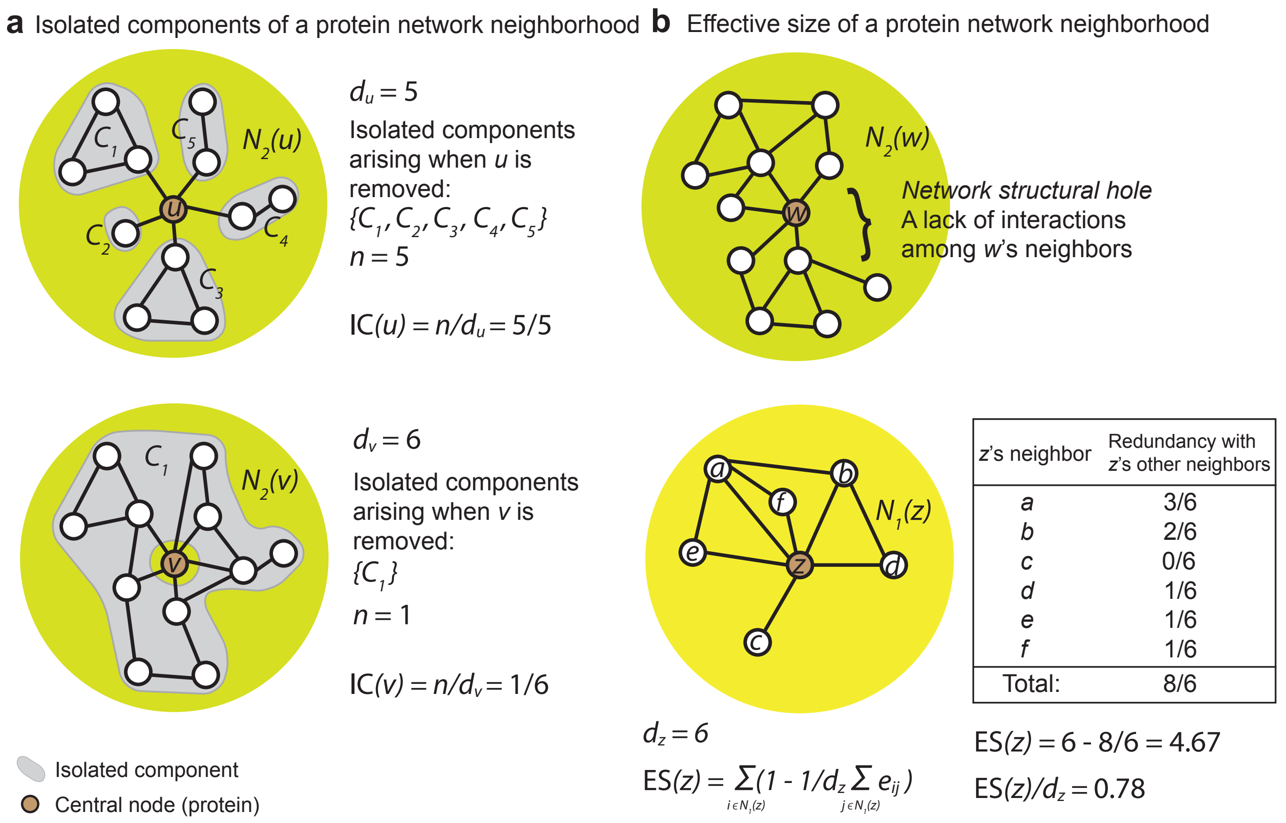


**Figure S5: Characterization of protein network neighborhoods.** Shown are network neighborhoods (Figure S4) of four hypothetical proteins, $u$, $v$, $w$, and $z$. The neighborhoods are characterized through two network metrics as follows (see also Section S7). **(a)** Isolated components metric (IC) is defined as the number of isolated components $n$ that arise when the central node is removed from the neighborhood. The metric is normalized by degree of the central node (*i.e.*, $d_u$, $d_v$) such that its maximum value is 1 and that a higher value indicates a greater fragmentation of the neighborhood. **(b)** Effective size metric (ES) captures the bridging potential of the central node, *i.e.*, the "true" size of the node's neighborhood absent of redundant neighbors (*49–51*). Taking a central node $w$ of the neighborhood, a redundant neighbor of $w$ is one that is also connected to other neighbors of $w$. This means that when $w$ is connected to non-redundant neighbors, $w$ sits on a "bridge" between those separate areas of the local network neighborhood that are known as network structural holes (*49*). The ES metric is mathematically defined as the sum of the non-redundant portion of the central node's connections over all the neighbors $N_1$. Shown is an example that illustrates computation of ES for node $z$.

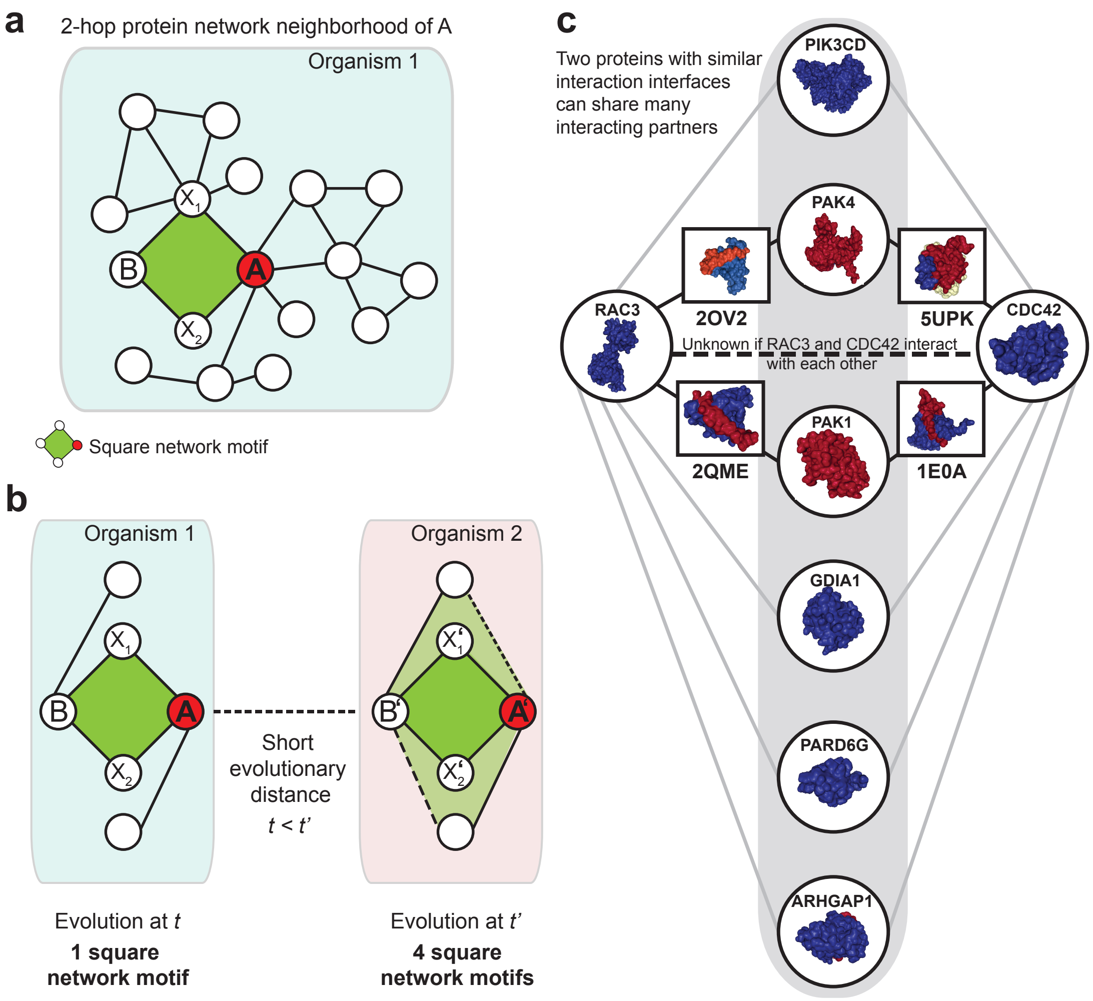


**Figure S6: Square network motifs of protein-protein interactions.** **(a)** Shown is a 2-hop protein network neighborhood (Figure S4) of protein $A$ in the interactome of Organism 1. Highlighted (in green) is an instance of a square network motif on nodes $A$, $X_1$, $X_2$, and $B$. **(b)** An illustration of the positive rate of change in the number of square network motifs (see Figure 4 in the main text for results on the protein-protein interaction dataset). There is one instance of a square network motif in the network neighborhood of protein $A$ in Organism 1. However, there are four instances of a square network motif in the network neighborhood of protein $A'$, an evolutionarily older ortholog of $A$ that is found in Organism 2. These additional instances are due to interaction rewiring (*i.e.*, two rewired/new PPIs/edges in Organism 2 are shown as dashed lines). **(c)** 3D structural illustration of our finding that proteins in evolutionarily older species have on average more square network motifs than proteins in evolutionarily younger species (see main text). To illustrate our finding with existing 3D structural data, we selected two human proteins from the Protein Data Bank (PDB) (*75*), RAC3 and CDC42, interacting with some of their partners through the same shared interface. While these two proteins are not known to interact with each other, we expect them to share some additional interacting partners, interacting with the same shared interface. Physical PPIs often require complementary interfaces. As a result, RAC3 and CDC42 with similar interfaces share many neighbors. Yet, it is not known if RAC3 and CDC42 directly interact with each other. Instead, additional interaction partner of RAC3 (protein PAK4) is also shared with protein CDC42 (in red). Besides PAK4, CDC42 has an additional interacting partner, PAK1, that potentially interact with RAC3 through the same interface. This detailed example illustrates that our finding on the positive rate of change in the number od square motifs agrees with structural and evolutionary arguments (*10, 76, 77*).

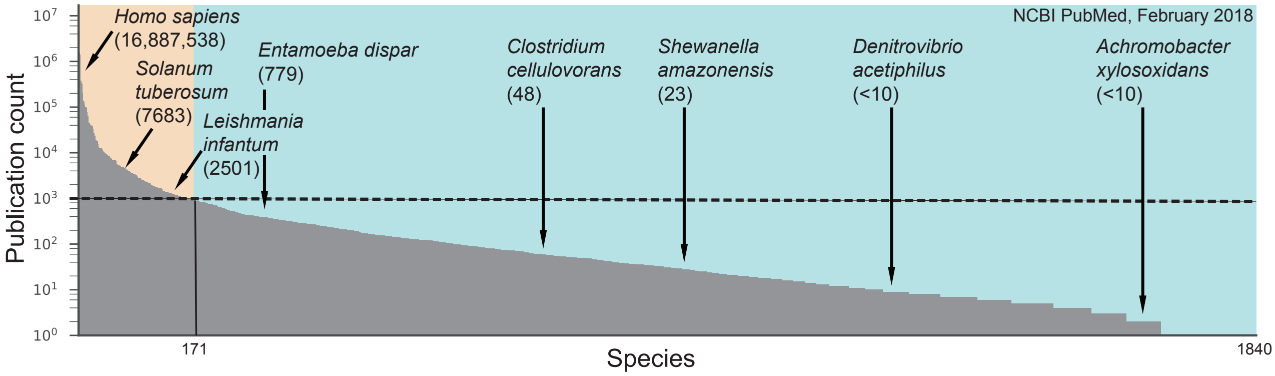


**Figure S7: Publication bias towards model organisms and highly studied species.** Shown is the number of publications in the NCBI Pubmed database (https://www.ncbi.nlm.nih.gov/pubmed) for each of 1,840 species. The publication data was obtained in February 2018. We used BioPython (*78*) and Entrez Programming Utilities (http://www.ncbi.nlm.nih.gov/books/NBK25501) to access the NCBI over programming interface and then used the NCBI's ESearch utility (https://eutils.ncbi.nlm.nih.gov/entrez/eutils/esearch.fcgi) to search and retrieve primary publication IDs and term translations based on species' names (*i.e.*, db='pubmed', term='species_name' or term='species_name [MeSH Terms]'). The plot reveals a substantial publication bias towards prominent model organisms and other highly studied species, suggesting that current protein-protein interaction data might be prone to notable selection and investigative biases (*1, 11*); hence we perform all analyses using only data from species that have at least 1,000 publications (dashed line; see also Section S1 and Table S5).

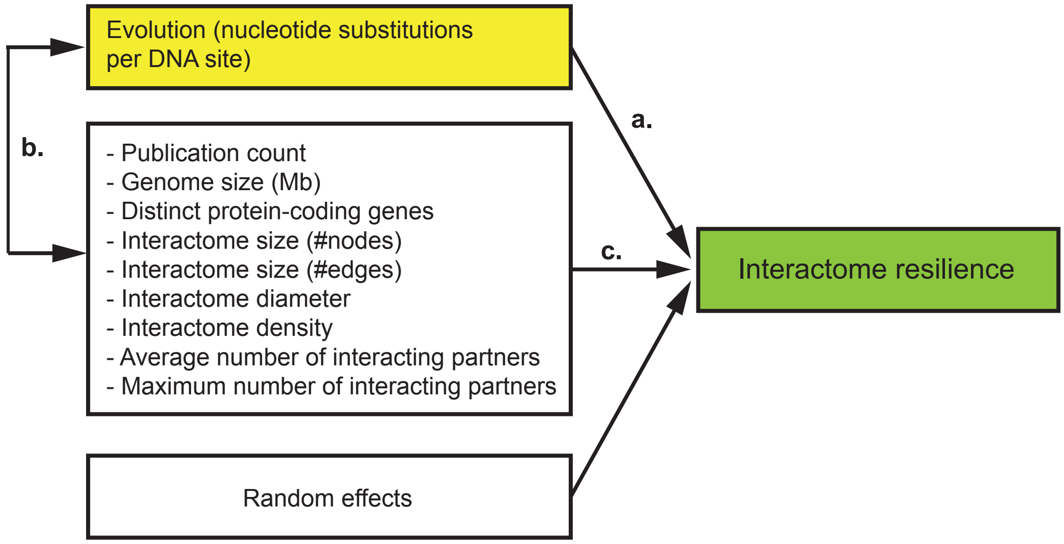


**Figure S8: Causal model for alternative hypotheses to explain the relationship between evolution and interactome resilience.** One hypothesis, represented by arrow a, is that interactomes become more resilient during evolution, indicating that a species' position in the tree of life is predictive of how robust the species' interactome is to network failures. Secondary hypotheses, represented by arrows b and c, are that non-biological (*e.g.*, the amount of research on a given species, the number of documented protein-protein interactions in a species) and other biological factors (*e.g.*, genome size, the number of a species' protein-coding genes) have a greater effect on a species' interactome and therefore better explain the resilience of the interactome. The secondary hypotheses can be rejected because these non-biological and biological factors cannot explain the observed relationship between evolution and interactome resilience (Table S1), indicating that our main results are not direct effects of various properties of species' genomes and interactome networks (Section S8).

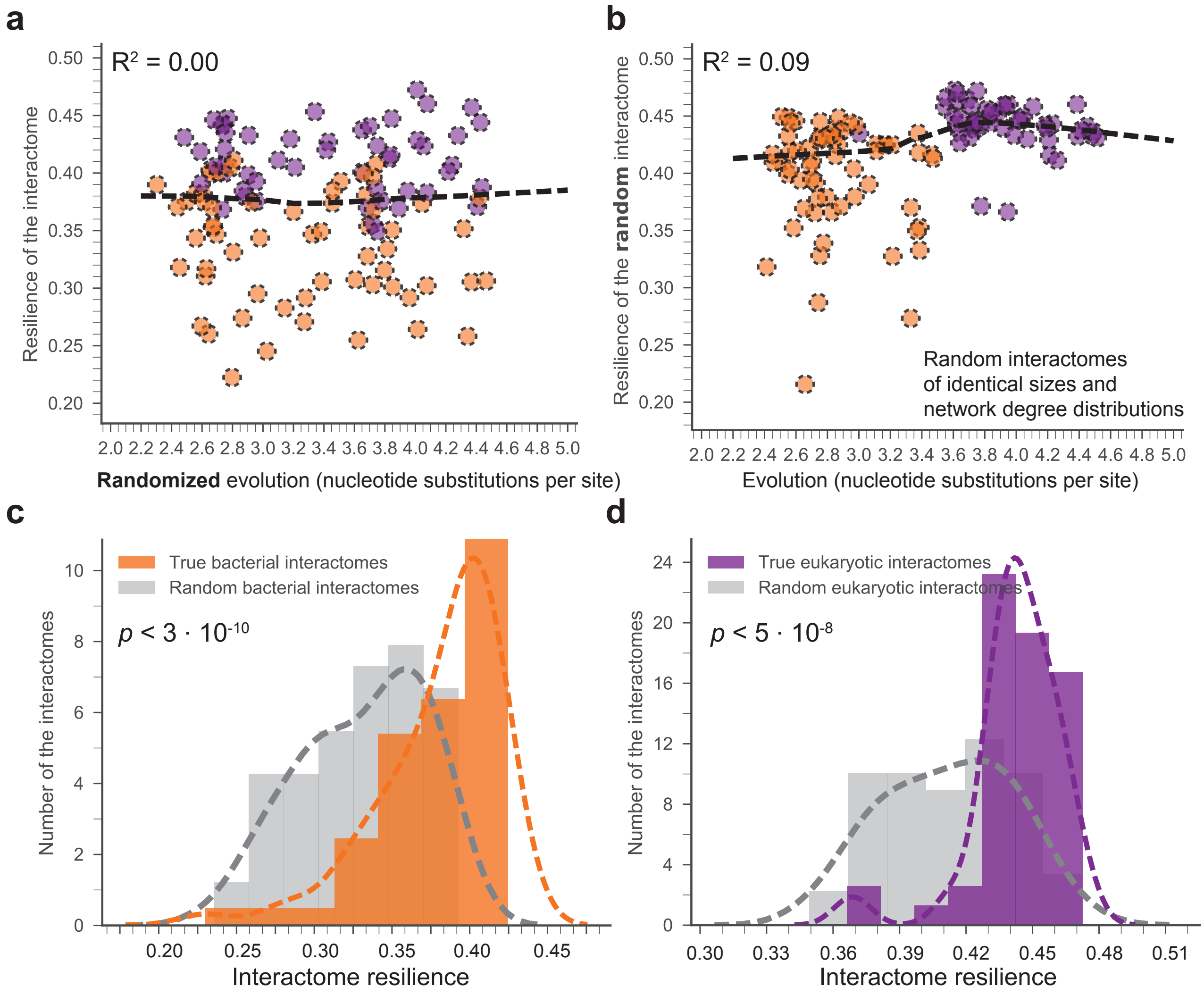


**Figure S9: Relationship between evolution and interactome resilience under random expectation.** We use two complementary null models (in **(a)** and **(b)**) to evaluate the statistical significance of the observed relationship between evolution and interactome resilience ($R^2 = 0.36$; see Figure 1 in the main text). **(a)** For a statistical evaluation of the observed relationship, we use a permutation model with the null hypothesis that evolution (*i.e.*, the amount of nucleotide substitutions per site) of a species is randomly drawn from the space of all possible nucleotide substitution rates. Comparing the observed relationship with random expectation, we find no significant association between *randomized* evolution and interactome resilience ($R^2 = 0.00$). **(b)** In a complementary random control, we use a configuration network model (see Section S7 for full details) with the null hypothesis that interactome of a species is randomly drawn from the space of all networks with identical sizes and degree distributions as the true species' interactome. Comparing the observed relationship with random expectation, we again find that the observed relationship cannot be explained solely by network size and degree distribution ($R^2 = 0.09$ for *random* interactomes vs. $R^2 = 0.36$ for true interactomes). **(c)** The distribution of the resilience for bacterial interactomes. Resilience of naturally occurring interactomes is significantly shifted to higher values compared to the random expectation ($p$ value $< 3 \cdot 10^{-10}$; denotes the significance of the difference of distributions using a non-parametric two-sided Mann-Whitney rank test). The expected distribution for random interactomes of identical sizes and degree distributions is shown in grey; the lines represent Gaussian kernel density estimates. **(d)** The distribution of the resilience for eukaryotic interactomes. Again, resilience of naturally occurring interactomes is significantly shifted to higher values compared to the random expectation ($p$ value $< 5 \cdot 10^{-8}$; denotes the significance of the difference of distributions using a non-parametric two-sided Mann-Whitney rank test).

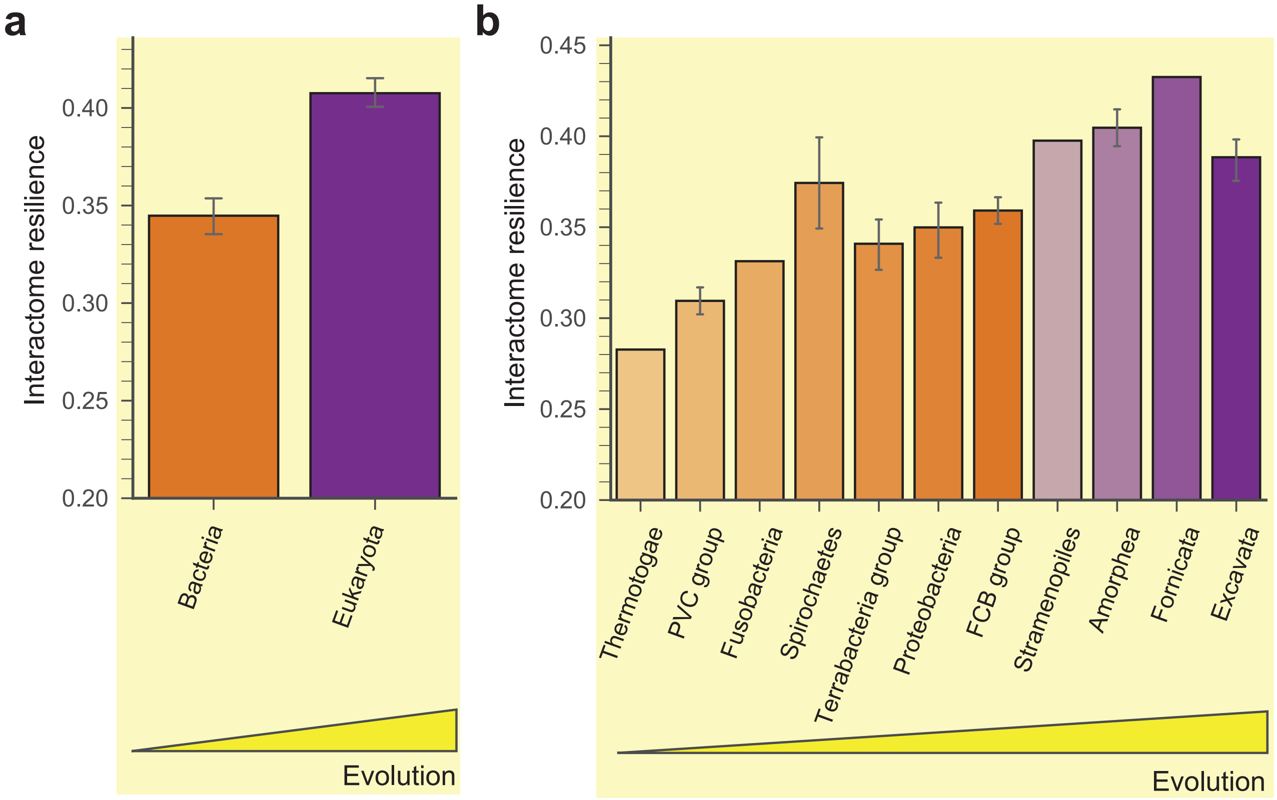


**Figure S10: Interactome resilience for species from the same taxonomic groups.** **(a)** Species from the same domain have more similar interactome resilience than species from different domains ($p$ value $= 6 \cdot 10^{-11}$). Error bars indicate 95% bootstrap confidence interval. **(b)** Species from the same taxonomic group (*i.e.*, supergroups or phyla) have similar interactome resilience. Furthermore, species in taxonomic groups with more nucleotide substitutions per site tend to have more resilient interactome. This observation is consistent with the main finding that a greater amount of genetic change is associated with a more resilient interactome structure. Taxonomic groups are defined based on the NCBI Taxonomy database (*17*) and the supergroups/phyla deliniated by Hug *et al.* (*13*) (Section S2). To obtain a higher resolution of the lineages, this analysis considers species with at least 500 publications in the NCBI Pubmed database (this gives 246 species; see also Figure S7). The bars, representing taxonomic groups, are ordered by the median evolution of species in each group. Colors indicate the assignment of taxonomic groups to domains; error bars indicate 95% bootstrap confidence interval.

#### Supplementary Tables

**Table S1: Analysis of possible confounding factors for interactome resilience.** We investigate the relationship between evolution ($E$) and interactome resilience ($R$) after removing the effects of possible confounding factors ($Z$). We perform a partial correlation analysis (Section S8) to investigate the extent to which our results could be explained by other biological and non-biological factors, such as genome size and the number of documented protein-protein interactions. To this aim, we quantify the relationship between evolution and interactome resilience while a particular confounding factor is held constant. The parametric and nonparametric partial measures of association indicate a significant correlation between evolution and interactome resilience that does not depend on and cannot be explained by these biological and non-biological factors. As with the standard (rank) correlation coefficient, a value (*i.e.*, $r_{ER|Z}$ and $\rho_{ER|Z}$) of +1 indicates a perfect positive linear relationship, a value of -1 indicates a perfect negative linear relationship, and a value of 0 indicates no linear relationship after removing the effect of a possible confounding factor.

| Possible confounding factor ($Z$) | Parametric partial measure of association | | | Nonparametric partial measure of association | |
|---|---|---|---|---|---|
| | $r_{ER\|Z}$ | $R^2_{ER\|Z}$ | $p$ value | $\rho_{ER\|Z}$ | $p$ value |
| Publication count | 0.545 | 0.30 | $5.15 \cdot 10^{-10}$ | 0.519 | $4.73 \cdot 10^{-9}$ |
| Genome size (Mb) | 0.472 | 0.22 | $2.19 \cdot 10^{-7}$ | 0.289 | $3.06 \cdot 10^{-2}$ |
| Distinct protein-coding genes | 0.359 | 0.13 | $1.02 \cdot 10^{-4}$ | 0.185 | $4.01 \cdot 10^{-2}$ |
| Interactome size (#nodes) | 0.496 | 0.25 | $2.49 \cdot 10^{-8}$ | 0.426 | $2.81 \cdot 10^{-6}$ |
| Interactome size (#edges) | 0.507 | 0.26 | $1.16 \cdot 10^{-8}$ | 0.390 | $2.09 \cdot 10^{-5}$ |
| Interactome diameter | 0.497 | 0.25 | $2.46 \cdot 10^{-8}$ | 0.484 | $6.24 \cdot 10^{-8}$ |
| Interactome density | 0.547 | 0.30 | $4.14 \cdot 10^{-10}$ | 0.506 | $1.18 \cdot 10^{-8}$ |
| Average number of interacting partners (avg. node degree) | 0.434 | 0.19 | $1.69 \cdot 10^{-6}$ | 0.178 | $6.03 \cdot 10^{-5}$ |
| Maximum number of interacting partners (max. node degree) | 0.500 | 0.25 | $1.89 \cdot 10^{-8}$ | 0.311 | $8.23 \cdot 10^{-4}$ |

**Table S2: Quality-controlled analysis of interactome data generated by yeast two-hybrid (Y2H) assays.** The protein-protein interaction dataset is prone to investigative biases (Section S1). We explore how our results are affected when only unbiased high-throughput data are used to quantify interactome resilience (Section S8). The table shows how interactome resilience of *H. sapiens* and *S. cerevisiae* relate to each other when only high-throughput interactions from various yeast two-hybrid assays are used instead of the full species' interactomes. The symbol '+' indicates the relationship between *H. sapiens* and *S. cerevisiae* persists also in the high-throughput Y2H data. The symbol '–' indicates the relationship in the high-throughput Y2H data is reversed relative to the relationship observed in the full species' interactomes. In other words, symbol '–' indicates an inconsistency: *S. cerevisiae* has higher interactome resilience than *H. sapiens* according to the full data but it has lower interactome resilience than *H. sapiens* when only the high-throughput Y2H data are used. Based on these results, we conclude that within the limitations imposed by the current protein-protein interaction data the interactome resilience continues to exist in unbiased high-throughput data (*i.e.*, in 17/20 dataset combinations) and that the values of interactome resilience correlate strongly between different combinations of data sources.

| High-throughput Y2H assays | | *H. sapiens* | | | |
|---|---|---|---|---|---|
| | | Rual *et al.* (*62*) | Stelzl *et al.* (*72*) | Venkatesan *et al.* (*64*) | Rolland *et al.* (*11*) |
| *S. cerevisiae* | Ito *et al.* (*70*) | + | + | + | + |
| | Krogan *et al.* (*71*) | + | + | + | + |
| | Yu *et al.* (*31*) | + | + | – | + |
| | Sahasranaman *et al.* (*73*) | – | + | + | + |
| | Porter *et al.* (*74*) | – | + | + | + |

**Table S3: Resilience of species' interactomes to network failure of essential protein-coding genes.** Essential protein-coding genes are indispensable for survival of an organism and are therefore considered a foundation of life. For example, in *S. cerevisiae*, these are genes whose mutant organisms are not viable ("inviability" phenotype is represented by APO:0000112 ontological term in the Yeast Phenotype Ontology, https://www.yeastgenome.org/ontology/phenotype/ypo), meaning that a mutant organism is not able to grow under standard growth conditions. This means that visible yeast colonies are not formed from single cells on plates rich with nutrients under normal atmospheric conditions (*79*). As another example, essential genes in bacteria constitute a minimal genome and encode proteins with essential functions, such as phosphate transport, that play key roles in organism survival (*80*). We obtain information on essential genes for six species. For each species, we quantify the resilience of species' interactome with respect to failure of essential genes in that species (Section S5). We find that interactomes are significantly less resilient to failures of essential genes than to failures of random genes/proteins ($p$ value $< 1 \cdot 10^{-4}$; permutation test). This finding is consistent across species and demonstrates that interactomes have a topological structure that is error-tolerant but extremely vulnerable to targeted attacks on essential genes. When essential genes are targeted the interactomes become rapidly fragmented and break into many small isolated components. This decrease in resilience provides evidence for the topological instability of interactomes to targeted attacks on essential genes. See Section S5.4 for a detailed discussion. 'Reference' indicates the source of gene essentiality information; '#essential' shows the number essential protein-coding genes; a lower value in 'Essential' column indicates a greater vulnerability of the interactome to attacks on essential genes.

| | | | Node removal strategy | | |
|---|---|---|---|---|---|
| Species | Reference | #essential | Random | Essential | $p$ value |
| *S. cerevisiae* | Cherry *et al.* (*79*), Giaver *et al.* (*81*) | 1,110 | 0.471 | 0.132 | $< 1 \cdot 10^{-4}$ |
| *H. sapiens* | Luo *et al.* (*82*), Wang *et al.* (*83*), Hart *et al.* (*84*) | 8,256 | 0.461 | 0.102 | $< 1 \cdot 10^{-4}$ |
| *M. musculus* | Luo *et al.* (*82*), Dickinson *et al.* (*85*) | 2,443 | 0.447 | 0.156 | $< 1 \cdot 10^{-4}$ |
| *D. melanogaster* | Luo *et al.* (*82*) | 339 | 0.424 | 0.169 | $< 1 \cdot 10^{-4}$ |
| *C. elegans* | Luo *et al.* (*82*), Kamath *et al.* (*86*) | 294 | 0.421 | 0.214 | $< 1 \cdot 10^{-4}$ |
| *A. thaliana* | Luo *et al.* (*82*), Meinke *et al.* (*87*) | 356 | 0.430 | 0.187 | $< 1 \cdot 10^{-4}$ |

**Table S4: Summary of dataset statistics for species and their genomes.** Summary of genome statistics for 114 species. Species are ordered by the number of protein-coding genes. Taxon ID refers to taxon identifiers of species based on the NCBI Taxonomy database (https://www.ncbi.nlm.nih.gov/taxonomy). Assembly accession refers to GenBank assembly accession identifiers based on the NCBI Assembly database (https://www.ncbi.nlm.nih.gov/assembly). Table continued on next page.

| Species | Taxon ID | Assembly accession | Status | Size (Mb) | #genes (↓) |
|---|---|---|---|---|---|
| *Glycine max* | 3847 | GCA.000004515.3 | Chromosome | 978.972 | 46,824 |
| *Oryza sativa Indica* | 39946 | GCA.001305255.1 | Chromosome | 352.121 | 40,745 |
| *Zea mays* | 4577 | GCA.000005005.6 | Chromosome | 2135.080 | 39,498 |
| *Oryza sativa Japonica* | 39947 | GCA.001433935.1 | Chromosome | 374.423 | 35,825 |
| *Solanum lycopersicum* | 4081 | GCA.000188115.2 | Chromosome | 823.786 | 34,675 |
| *Sorghum bicolor* | 4558 | GCA.000003195.3 | Chromosome | 709.345 | 34,496 |
| *Vitis vinifera* | 29760 | GCA.000003745.2 | Chromosome | 486.197 | 29,927 |
| *Arabidopsis thaliana* | 3702 | GCA.000001735.1 | Chromosome | 119.668 | 27,416 |
| *Danio rerio* | 7955 | GCA.000002035.4 | Chromosome | 1679.200 | 26,163 |
| *Rattus norvegicus* | 10116 | GCA.000001895.4 | Chromosome | 2870.180 | 22,941 |
| *Mus musculus* | 10090 | GCA.000001635.8 | Chromosome | 2818.970 | 22,668 |
| *Macaca mulatta* | 9544 | GCA.000772875.3 | Chromosome | 3236.220 | 21,905 |
| *Sus scrofa* | 9823 | GCA.000003025.6 | Chromosome | 2501.910 | 21,630 |
| *Oreochromis niloticus* | 8128 | GCA.001858045.2 | Chromosome | 1009.860 | 21,437 |
| *Callithrix jacchus* | 9483 | GCA.000004665.1 | Chromosome | 2914.960 | 20,993 |
| *Caenorhabditis elegans* | 6239 | GCA.000002985.3 | Complete Genome | 100.286 | 20,517 |
| *Homo sapiens* | 9606 | GCA.000001405.26 | Chromosome | 3253.850 | 20,457 |
| *Equus caballus* | 9796 | GCA.002863925.1 | Chromosome | 2506.970 | 20,449 |
| *Bos taurus* | 9913 | GCA.000003055.5 | Chromosome | 2670.140 | 19,994 |
| *Oryzias latipes* | 8090 | GCA.002234675.1 | Chromosome | 734.057 | 19,686 |
| *Felis catus* | 9685 | GCA.000181335.4 | Chromosome | 2521.860 | 19,493 |
| *Oryctolagus cuniculus* | 9986 | GCA.000003625.1 | Chromosome | 2737.460 | 19,018 |
| *Pan troglodytes* | 9598 | GCA.000001515.5 | Chromosome | 3231.170 | 18,759 |
| *Gallus gallus* | 9031 | GCA.000002315.3 | Chromosome | 1230.260 | 16,736 |
| *Ciona intestinalis* | 7719 | GCA.000224145.2 | Chromosome | 115.227 | 16,658 |
| *Tribolium castaneum* | 7070 | GCA.000002335.3 | Chromosome | 165.944 | 16,524 |
| *Aedes aegypti* | 7159 | GCA.002204515.1 | Chromosome | 1278.730 | 15,998 |
| *Drosophila melanogaster* | 7227 | GCA.000001215.4 | Chromosome | 143.726 | 13,937 |
| *Schistosoma mansoni* | 6183 | GCA.000237925.2 | Chromosome | 364.538 | 11,770 |
| *Apis mellifera* | 7460 | GCA.000002195.1 | Chromosome | 250.287 | 10,694 |
| *Leishmania braziliensis* | 420245 | GCA.000002845.2 | Chromosome | 32.069 | 8,160 |
| *Leishmania infantum* | 435258 | GCA.000002875.2 | Chromosome | 32.122 | 8,150 |
| *Leishmania donovani* | 5661 | GCA.000227135.2 | Chromosome | 32.445 | 8,032 |

| | | | | | |
|---|---|---|---|---|---|
| *Streptomyces coelicolor* | 100226 | GCA.000203835.1 | Complete Genome | 9.055 | 7,768 |
| *Streptomyces griseus* | 455632 | GCA.000010605.1 | Complete Genome | 8.546 | 7,136 |
| *Mycobacterium smegmatis* | 246196 | GCA.000015005.1 | Complete Genome | 6.988 | 6,717 |
| *Saccharomyces cerevisiae* | 4932 | GCA.001051215.1 | Complete Genome | 12.086 | 6,692 |
| *Cryptococcus neoformans B* | 283643 | GCA.000149385.1 | Chromosome | 19.700 | 6,578 |
| *Streptomyces sp. SirexAAE* | 862751 | GCA.000177195.2 | Complete Genome | 7.414 | 6,357 |
| *Microcystis aeruginosa* | 449447 | GCA.000010625.1 | Complete Genome | 5.843 | 6,312 |
| *Agrobacterium radiobacter* | 311403 | GCA.000016265.1 | Complete Genome | 7.273 | 6,107 |
| *Corynebacterium glutamicum* | 196627 | GCA.000011325.1 | Complete Genome | 3.309 | 6,050 |
| *Vibrio harveyi* | 338187 | GCA.000017705.1 | Complete Genome | 6.058 | 5,921 |
| *Lactobacillus rhamnosus* | 568703 | GCA.000026505.1 | Complete Genome | 3.010 | 5,747 |
| *Burkholderia pseudomallei* | 272560 | GCA.000011545.1 | Complete Genome | 7.248 | 5,728 |
| *Pseudomonas aeruginosa* | 208964 | GCA.000006765.1 | Complete Genome | 6.264 | 5,571 |
| *Mycobacterium gilvum* | 350054 | GCA.000016365.1 | Complete Genome | 5.983 | 5,241 |
| *Schizosaccharomyces pombe* | 4896 | GCA.000002945.2 | Chromosome | 12.591 | 5,144 |
| *Bacillus megaterium* | 545693 | GCA.000025825.1 | Complete Genome | 5.523 | 5,116 |
| *Pseudomonas syringae syringae* | 205918 | GCA.000012245.1 | Complete Genome | 6.094 | 5,089 |
| *Plasmodium vivax* | 5855 | GCA.000002415.2 | Chromosome | 27.014 | 5,050 |
| *Enterobacter aerogenes* | 1028307 | GCA.000215745.1 | Complete Genome | 5.280 | 4,912 |
| *Plasmodium berghei* | 5821 | GCA.900044335.1 | Chromosome | 18.811 | 4,881 |
| *Vibrio parahaemolyticus* | 223926 | GCA.000196095.1 | Complete Genome | 5.166 | 4,832 |
| *Klebsiella pneumoniae* | 272620 | GCA.000016305.1 | Complete Genome | 5.695 | 4,776 |
| *Shigella flexneri* | 198214 | GCA.000006925.2 | Complete Genome | 4.829 | 4,439 |
| *Mycobacterium avium* | 262316 | GCA.000007865.1 | Complete Genome | 4.830 | 4,350 |
| *Bacillus subtilis 168* | 224308 | GCA.000009045.1 | Complete Genome | 4.216 | 4,280 |
| *Aeromonas hydrophila* | 380703 | GCA.000014805.1 | Complete Genome | 4.744 | 4,121 |
| *Stenotrophomonas maltophilia R5513* | 391008 | GCA.000020665.1 | Complete Genome | 4.574 | 4,039 |
| *Mycobacterium tuberculosis H37Rv* | 83332 | GCA.000195955.2 | Complete Genome | 4.412 | 4,003 |
| *Yersinia enterocolitica* | 393305 | GCA.000009345.1 | Complete Genome | 4.684 | 3,978 |
| *Cryptosporidium parvum* | 353152 | GCA.000165345.1 | Chromosome | 9.102 | 3,805 |
| *Rhodospirillum rubrum* | 269796 | GCA.000013085.1 | Complete Genome | 4.407 | 3,788 |
| *Vibrio fischeri* | 312309 | GCA.000011805.1 | Complete Genome | 4.274 | 3,760 |
| *Vibrio anguillarum* | 882102 | GCA.000217675.1 | Complete Genome | 4.052 | 3,732 |
| *Clostridium difficile* | 272563 | GCA.000009205.1 | Complete Genome | 4.298 | 3,728 |
| *Enterobacter cloacae NCTC9394* | 718254 | GCA.000210775.1 | Chromosome | 4.909 | 3,725 |
| *Proteus mirabilis* | 529507 | GCA.000069965.1 | Complete Genome | 4.100 | 3,607 |
| *Rhodobacter capsulatus* | 272942 | GCA.000021865.1 | Complete Genome | 3.872 | 3,493 |
| *Bordetella pertussis* | 257313 | GCA.000195715.1 | Complete Genome | 4.086 | 3,436 |
| *Sinorhizobium meliloti* | 266834 | GCA.000006965.1 | Complete Genome | 6.692 | 3,359 |
| *Enterococcus faecalis* | 226185 | GCA.000007785.1 | Complete Genome | 3.360 | 3,112 |
| *Rhodobacter sphaeroides ATCC17025* | 349102 | GCA.000016405.1 | Complete Genome | 4.557 | 3,111 |

| | | | | | |
|---|---|---|---|---|---|
| *Brucella melitensis* | 224914 | GCA.000007125.1 | Complete Genome | 3.295 | 3,083 |
| *Lactobacillus casei* | 543734 | GCA.000026485.1 | Complete Genome | 3.079 | 3,015 |
| *Lactobacillus plantarum* | 220668 | GCA.000203855.3 | Complete Genome | 3.349 | 3,007 |
| *Brucella abortus* | 430066 | GCA.000018725.1 | Complete Genome | 3.284 | 3,000 |
| *Listeria innocua* | 272626 | GCA.000195795.1 | Complete Genome | 3.093 | 2,968 |
| *Synechococcus sp. JA23Ba213* | 321332 | GCA.000013225.1 | Complete Genome | 3.047 | 2,862 |
| *Thiobacillus denitrificans* | 292415 | GCA.000012745.1 | Complete Genome | 2.910 | 2,827 |
| *Sulfolobus tokodaii* | 273063 | GCA.000011205.1 | Complete Genome | 2.695 | 2,826 |
| *Sulfolobus islandicus* | 930945 | GCA.000189555.1 | Complete Genome | 2.523 | 2,644 |
| *Flavobacteriaceae bacterium 351910* | 531844 | GCA.000023725.1 | Complete Genome | 2.768 | 2,534 |
| *Aggregatibacter actinomycetemcomitans* | 694569 | GCA.000163615.3 | Complete Genome | 2.309 | 2,432 |
| *Corynebacterium diphtheriae* | 698964 | GCA.000255275.1 | Complete Genome | 2.531 | 2,322 |
| *Lactococcus lactis lactis* | 272623 | GCA.000006865.1 | Complete Genome | 2.366 | 2,321 |
| *Streptococcus suis* | 391295 | GCA.000014305.1 | Complete Genome | 2.096 | 2,186 |
| *Streptococcus agalactiae* | 211110 | GCA.000196055.1 | Complete Genome | 2.211 | 2,094 |
| *Fusobacterium nucleatum nucleatum* | 190304 | GCA.000007325.1 | Complete Genome | 2.175 | 2,063 |
| *Neisseria meningitidis* | 122586 | GCA.000008805.1 | Complete Genome | 2.272 | 2,063 |
| *Pasteurella multocida* | 272843 | GCA.000006825.1 | Complete Genome | 2.257 | 2,012 |
| *Francisella sp. TX077308* | 573569 | GCA.000219045.1 | Complete Genome | 2.036 | 1,976 |
| *Chlamydophila psittaci* | 331636 | GCA.000204255.1 | Complete Genome | 1.179 | 1,970 |
| *Streptococcus mutans* | 210007 | GCA.000007465.2 | Complete Genome | 2.033 | 1,960 |
| *Thermotoga maritima* | 243274 | GCA.000008545.1 | Complete Genome | 1.861 | 1,858 |
| *Coxiella burnetii* | 227377 | GCA.000007765.2 | Complete Genome | 2.033 | 1,817 |
| *Streptococcus pyogenes* | 160490 | GCA.000006785.2 | Complete Genome | 1.852 | 1,696 |
| *Borrelia afzelii* | 390236 | GCA.000222835.1 | Complete Genome | 1.404 | 1,675 |
| *Haemophilus influenzae* | 71421 | GCA.000027305.1 | Complete Genome | 1.830 | 1,657 |
| *Mycobacterium leprae* | 272631 | GCA.000195855.1 | Complete Genome | 3.268 | 1,605 |
| *Francisella tularensis tularensis* | 177416 | GCA.000008985.1 | Complete Genome | 1.893 | 1,604 |
| *Helicobacter pylori SouthAfrica7* | 907239 | GCA.000185245.1 | Complete Genome | 1.680 | 1,543 |
| *Bartonella henselae* | 283166 | GCA.000046705.1 | Complete Genome | 1.931 | 1,488 |
| *Anaplasma phagocytophilum* | 212042 | GCA.000013125.1 | Complete Genome | 1.471 | 1,264 |
| *Orientia tsutsugamushi* | 357244 | GCA.000063545.1 | Complete Genome | 2.127 | 1,182 |
| *Treponema pallidum* | 243276 | GCA.000410535.2 | Chromosome | 1.140 | 1,036 |
| *Anaplasma marginale StMaries* | 234826 | GCA.000011945.1 | Complete Genome | 1.198 | 948 |
| *Chlamydia trachomatis* | 272561 | GCA.000008725.1 | Complete Genome | 1.043 | 895 |
| *Borrelia garinii* | 290434 | GCA.000196215.1 | Complete Genome | 0.987 | 829 |
| *Borrelia burgdorferi* | 224326 | GCA.000008685.2 | Complete Genome | 1.521 | 753 |
| *Mycoplasma putrefaciens* | 743965 | GCA.000224105.1 | Complete Genome | 0.833 | 650 |
| *Mycoplasma pneumoniae* | 722438 | GCA.000143945.1 | Complete Genome | 0.811 | 629 |
| *Ureaplasma parvum* | 273119 | GCA.000006625.1 | Complete Genome | 0.752 | 614 |

**Table S5: Summary of interactome resilience and dataset statistics for species and interactomes.** Summary of dataset statistics for the 171 species with more than 1000 publications in the NCBI Pubmed (Figure S7; Section S1). Species are ordered by the interactome resilience (Section S5). Domain and group refer to taxonomic information of species (Section S2) based on the NCBI Taxonomy database (https://www.ncbi.nlm.nih.gov/taxonomy) and Hug *et al.* (*13*). Table continued on next page.

| Species | Domain | Group | Pub. count | #nodes | #edges | Resilience (↓) |
|---|---|---|---|---|---|---|
| *Saccharomyces cerevisiae* | Eukaryota | Opisthokonta | 96,928 | 6,011 | 207,622 | 0.471 |
| *Homo sapiens* | Eukaryota | Opisthokonta | 16,887,538 | 16,439 | 440,135 | 0.461 |
| *Glycine max* | Eukaryota | Viridiplantae | 19,004 | 5,785 | 89,538 | 0.457 |
| *Oryza sativa Japonica* | Eukaryota | Viridiplantae | 1,639 | 1,787 | 54,247 | 0.453 |
| *Bos taurus* | Eukaryota | Opisthokonta | 328,217 | 8,615 | 276,128 | 0.453 |
| *Sus scrofa* | Eukaryota | Opisthokonta | 17,660 | 8,201 | 143,516 | 0.450 |
| *Callithrix jacchus* | Eukaryota | Opisthokonta | 3,486 | 327 | 946 | 0.448 |
| *Rattus norvegicus* | Eukaryota | Opisthokonta | 1,536,193 | 9,439 | 261,737 | 0.447 |
| *Mus musculus* | Eukaryota | Opisthokonta | 1,399,668 | 12,498 | 354,458 | 0.447 |
| *Oryctolagus cuniculus* | Eukaryota | Opisthokonta | 331,624 | 292 | 844 | 0.447 |
| *Magnaporthe grisea* | Eukaryota | Opisthokonta | 1,192 | 26 | 68 | 0.446 |
| *Oreochromis niloticus* | Eukaryota | Opisthokonta | 4,060 | 32 | 55 | 0.444 |
| *Aspergillus terreus* | Eukaryota | Opisthokonta | 1,396 | 45 | 138 | 0.444 |
| *Danio rerio* | Eukaryota | Opisthokonta | 23,531 | 7,377 | 145,449 | 0.443 |
| *Cavia porcellus* | Eukaryota | Opisthokonta | 138,133 | 165 | 268 | 0.441 |
| *Oryza sativa Indica* | Eukaryota | Viridiplantae | 1,597 | 105 | 224 | 0.441 |
| *Solanum lycopersicum* | Eukaryota | Viridiplantae | 10,259 | 3,601 | 32,470 | 0.440 |
| *Mustela putorius* | Eukaryota | Opisthokonta | 5,403 | 173 | 254 | 0.439 |
| *Oryzias latipes* | Eukaryota | Opisthokonta | 2,281 | 4,134 | 35,414 | 0.439 |
| *Physcomitrella patens* | Eukaryota | Viridiplantae | 1,050 | 3,190 | 31,790 | 0.438 |
| *Gallus gallus* | Eukaryota | Opisthokonta | 112,314 | 5,208 | 70,943 | 0.437 |
| *Sorghum bicolor* | Eukaryota | Viridiplantae | 1,775 | 3,503 | 26,577 | 0.435 |
| *Felis catus* | Eukaryota | Opisthokonta | 131,626 | 5,285 | 44,325 | 0.434 |
| *Ailuropoda melanoleuca* | Eukaryota | Opisthokonta | 2,111 | 5,034 | 41,328 | 0.433 |
| *Giardia lamblia* | Eukaryota | Fornicata | 2,474 | 238 | 453 | 0.433 |
| *Schizosaccharomyces pombe* | Eukaryota | Opisthokonta | 9,409 | 4,125 | 47,984 | 0.431 |
| *Arabidopsis thaliana* | Eukaryota | Viridiplantae | 45,738 | 11,256 | 167,771 | 0.430 |
| *Equus caballus* | Eukaryota | Opisthokonta | 64,851 | 5,395 | 45,074 | 0.430 |
| *Solanum tuberosum* | Eukaryota | Viridiplantae | 7,683 | 3,239 | 27,478 | 0.429 |
| *Vitis vinifera* | Eukaryota | Viridiplantae | 6,986 | 3,418 | 28,243 | 0.427 |
| *Hordeum vulgare* | Eukaryota | Viridiplantae | 8,527 | 50 | 152 | 0.424 |
| *Ixodes scapularis* | Eukaryota | Opisthokonta | 3,801 | 2,052 | 12,610 | 0.424 |
| *Drosophila melanogaster* | Eukaryota | Opisthokonta | 41,416 | 10,132 | 116,906 | 0.424 |

| | | | | | | |
|---|---|---|---|---|---|---|
| *Gorilla gorilla* | Eukaryota | Opisthokonta | 1,803 | 5,309 | 41,774 | 0.423 |
| *Trypanosoma cruzi* | Eukaryota | Euglenozoa | 10,781 | 1,267 | 5,258 | 0.422 |
| *Pan troglodytes* | Eukaryota | Opisthokonta | 8,981 | 5,166 | 39,642 | 0.422 |
| *Caenorhabditis elegans* | Eukaryota | Opisthokonta | 19,285 | 8,091 | 82,152 | 0.421 |
| *Dictyostelium discoideum* | Eukaryota | Amoebozoa | 7,004 | 2,065 | 24,430 | 0.420 |
| *Macaca mulatta* | Eukaryota | Opisthokonta | 38,314 | 5,003 | 37,070 | 0.419 |
| *Zea mays* | Eukaryota | Viridiplantae | 27,146 | 2,849 | 18,046 | 0.419 |
| *Aspergillus flavus* | Eukaryota | Opisthokonta | 2,348 | 1,668 | 9,044 | 0.416 |
| *Agrobacterium radiobacter* | Bacteria | Proteobacteria | 3,188 | 1,682 | 10,840 | 0.414 |
| *Streptomyces griseus* | Bacteria | Terrabacteria group | 1,208 | 1,432 | 9,882 | 0.413 |
| *Aspergillus fumigatus* | Eukaryota | Opisthokonta | 6,769 | 1,537 | 8,028 | 0.413 |
| *Tetrahymena thermophila* | Eukaryota | Alveolata | 1,147 | 1,093 | 5,323 | 0.411 |
| *Streptomyces coelicolor* | Bacteria | Terrabacteria group | 1,119 | 1,423 | 8,372 | 0.411 |
| *Coccidioides immitis* | Eukaryota | Opisthokonta | 1,290 | 1,193 | 4,438 | 0.410 |
| *Mycobacterium gilvum* | Bacteria | Terrabacteria group | 10,015 | 1,217 | 9,400 | 0.410 |
| *Bacillus subtilis 168* | Bacteria | Terrabacteria group | 1,263 | 1,494 | 5,896 | 0.410 |
| *Trichinella spiralis* | Eukaryota | Opisthokonta | 1,452 | 1,453 | 5,511 | 0.410 |
| *Bacillus megaterium* | Bacteria | Terrabacteria group | 2,780 | 1,300 | 6,355 | 0.408 |
| *Leishmania major* | Eukaryota | Euglenozoa | 2,736 | 729 | 2,537 | 0.407 |
| *Aspergillus niger* | Eukaryota | Opisthokonta | 5,103 | 1,279 | 5,391 | 0.407 |
| *Burkholderia pseudomallei* | Bacteria | Proteobacteria | 1,865 | 1,583 | 8,485 | 0.406 |
| *Leishmania donovani* | Eukaryota | Euglenozoa | 4,165 | 655 | 2,031 | 0.406 |
| *Leishmania braziliensis* | Eukaryota | Euglenozoa | 18,380 | 735 | 2,490 | 0.405 |
| *Aspergillus oryzae* | Eukaryota | Opisthokonta | 1,669 | 1,400 | 6,125 | 0.405 |
| *Sinorhizobium meliloti* | Bacteria | Proteobacteria | 1,586 | 956 | 4,182 | 0.405 |
| *Enterobacter aerogenes* | Bacteria | Proteobacteria | 1,008 | 1,529 | 7,351 | 0.403 |
| *Leishmania infantum* | Eukaryota | Euglenozoa | 2,501 | 725 | 2,347 | 0.401 |
| *Cryptococcus neoformans B* | Eukaryota | Opisthokonta | 1,773 | 1,134 | 3,986 | 0.401 |
| *Bordetella pertussis* | Bacteria | Proteobacteria | 4,971 | 1,004 | 5,716 | 0.401 |
| *Plasmodium berghei* | Eukaryota | Alveolata | 4,767 | 428 | 1,056 | 0.400 |
| *Borrelia garinii* | Bacteria | Spirochaetes | 6,808 | 186 | 408 | 0.399 |
| *Pseudomonas aeruginosa* | Bacteria | Proteobacteria | 38,172 | 1,529 | 8,273 | 0.399 |
| *Plasmodium vivax* | Eukaryota | Alveolata | 4,715 | 462 | 1,212 | 0.398 |
| *Streptomyces sp. SirexAAE* | Bacteria | Terrabacteria group | 3,637 | 1,397 | 7,732 | 0.396 |
| *Rhodobacter capsulatus* | Bacteria | Proteobacteria | 1,060 | 1,026 | 4,844 | 0.394 |
| *Stenotrophomonas maltophilia R5513* | Bacteria | Proteobacteria | 2,227 | 719 | 3,076 | 0.393 |
| *Trypanosoma brucei* | Eukaryota | Euglenozoa | 6,817 | 708 | 3,494 | 0.393 |
| *Vibrio parahaemolyticus* | Bacteria | Proteobacteria | 2,225 | 1,260 | 5,923 | 0.392 |
| *Pseudomonas syringae syringae* | Bacteria | Proteobacteria | 2,002 | 1,316 | 6,157 | 0.392 |
| *Penicillium chrysogenum* | Eukaryota | Opisthokonta | 1,231 | 1,597 | 7,358 | 0.392 |
| *Schistosoma mansoni* | Eukaryota | Opisthokonta | 9,526 | 958 | 3,520 | 0.391 |

| | | | | | | |
|---|---|---|---|---|---|---|
| *Pediculus humanus* | Eukaryota | Opisthokonta | 1,050 | 1,710 | 7,539 | 0.391 |
| *Aeromonas hydrophila* | Bacteria | Proteobacteria | 1,653 | 1,220 | 5,962 | 0.390 |
| *Plasmodium yoelii* | Eukaryota | Alveolata | 1,348 | 404 | 1,106 | 0.390 |
| *Treponema pallidum* | Bacteria | Spirochaetes | 3,696 | 664 | 1,625 | 0.389 |
| *Salmonella enterica RSK2980* | Bacteria | Proteobacteria | 45,567 | 1,226 | 5,006 | 0.388 |
| *Chlamydomonas reinhardtii* | Eukaryota | Viridiplantae | 3,429 | 1,449 | 8,207 | 0.388 |
| *Aedes aegypti* | Eukaryota | Opisthokonta | 8,483 | 1,994 | 8,913 | 0.387 |
| *Sulfolobus islandicus* | Archaea | TACK group | 2,145 | 546 | 2,788 | 0.385 |
| *Vibrio fischeri* | Bacteria | Proteobacteria | 1,097 | 1,074 | 4,514 | 0.385 |
| *Bombyx mori* | Eukaryota | Opisthokonta | 6,582 | 1,732 | 8,521 | 0.384 |
| *Tribolium castaneum* | Eukaryota | Opisthokonta | 1,343 | 1,734 | 8,073 | 0.384 |
| *Lactobacillus casei* | Bacteria | Terrabacteria group | 4,753 | 758 | 3,162 | 0.384 |
| *Plasmodium falciparum* | Eukaryota | Alveolata | 26,907 | 1,688 | 4,634 | 0.383 |
| *Rhodospirillum rubrum* | Bacteria | Proteobacteria | 1,365 | 1,092 | 5,502 | 0.382 |
| *Corynebacterium diphtheriae* | Bacteria | Terrabacteria group | 2,405 | 617 | 2,112 | 0.382 |
| *Enterococcus faecalis* | Bacteria | Terrabacteria group | 9,751 | 745 | 2,587 | 0.380 |
| *Neurospora crassa* | Eukaryota | Opisthokonta | 5,302 | 1,364 | 5,028 | 0.380 |
| *Rhodobacter sphaeroides ATCC17025* | Bacteria | Proteobacteria | 3,925 | 980 | 3,730 | 0.379 |
| *Thalassiobium sp. R2A62* | Bacteria | Proteobacteria | 205,044 | 517 | 821 | 0.378 |
| *Culex quinquefasciatus* | Eukaryota | Opisthokonta | 2,815 | 1,846 | 8,800 | 0.376 |
| *Anopheles gambiae* | Eukaryota | Opisthokonta | 12,397 | 1,808 | 7,214 | 0.376 |
| *Aggregatibacter actinomycetemcomitans* | Bacteria | Proteobacteria | 2,593 | 703 | 2,173 | 0.375 |
| *Vibrio anguillarum* | Bacteria | Proteobacteria | 1,204 | 1,049 | 4,945 | 0.375 |
| *Streptococcus suis* | Bacteria | Terrabacteria group | 1,047 | 561 | 1,713 | 0.375 |
| *Shigella flexneri* | Bacteria | Proteobacteria | 3,516 | 1,230 | 5,039 | 0.374 |
| *Proteus mirabilis* | Bacteria | Proteobacteria | 3,330 | 934 | 3,709 | 0.373 |
| *Kluyveromyces lactis* | Eukaryota | Opisthokonta | 1,451 | 1,437 | 5,636 | 0.371 |
| *Pasteurella multocida* | Bacteria | Proteobacteria | 1,806 | 706 | 2,380 | 0.370 |
| *Ciona intestinalis* | Eukaryota | Opisthokonta | 1,113 | 1,372 | 5,432 | 0.370 |
| *Francisella sp. TX077308* | Bacteria | Proteobacteria | 3,489 | 552 | 1,781 | 0.370 |
| *Apis mellifera* | Eukaryota | Opisthokonta | 4,156 | 1,561 | 6,583 | 0.369 |
| *Cryptosporidium parvum* | Eukaryota | Alveolata | 2,545 | 287 | 653 | 0.368 |
| *Lactobacillus plantarum* | Bacteria | Terrabacteria group | 1,889 | 774 | 2,885 | 0.368 |
| *Flavobacteriaceae bacterium 351910* | Bacteria | FCB group | 4,471 | 491 | 1,624 | 0.366 |
| *Borrelia afzelii* | Bacteria | Spirochaetes | 6,808 | 83 | 70 | 0.364 |
| *Mycobacterium leprae* | Bacteria | Terrabacteria group | 5,447 | 479 | 1,464 | 0.358 |
| *Lactococcus lactis lactis* | Bacteria | Terrabacteria group | 5,716 | 542 | 1,684 | 0.358 |
| *Klebsiella pneumoniae* | Bacteria | Proteobacteria | 12,105 | 773 | 1,707 | 0.357 |
| *Anaplasma phagocytophilum* | Bacteria | Proteobacteria | 1,005 | 279 | 642 | 0.356 |
| *Candida glabrata* | Eukaryota | Opisthokonta | 1,309 | 1,423 | 5,421 | 0.356 |
| *Neisseria meningitidis* | Bacteria | Proteobacteria | 8,868 | 565 | 1,614 | 0.355 |

| | | | | | | |
|---|---|---|---|---|---|---|
| *Streptococcus mutans* | Bacteria | Terrabacteria group | 8,264 | 521 | 1,589 | 0.354 |
| *Streptococcus sp. 73H25AP* | Bacteria | Terrabacteria group | 99,893 | 257 | 326 | 0.354 |
| *Aerococcus viridans* | Bacteria | Terrabacteria group | 77,845 | 399 | 587 | 0.353 |
| *Gasterosteus aculeatus* | Eukaryota | Opisthokonta | 1,040 | 45 | 57 | 0.353 |
| *Thiobacillus denitrificans* | Bacteria | Proteobacteria | 1,082 | 775 | 2,553 | 0.352 |
| *Coxiella burnetii* | Bacteria | Proteobacteria | 2,111 | 458 | 1,291 | 0.351 |
| *Clostridiales bacterium* | Bacteria | Terrabacteria group | 32,957 | 385 | 1,019 | 0.350 |
| *Toxoplasma gondii* | Eukaryota | Alveolata | 12,356 | 596 | 1,739 | 0.349 |
| *Lactobacillus rhamnosus* | Bacteria | Terrabacteria group | 1,041 | 969 | 2,911 | 0.349 |
| *Synechococcus sp. JA23Ba213* | Bacteria | Terrabacteria group | 1,292 | 686 | 2,034 | 0.347 |
| *Mycoplasma pneumoniae* | Bacteria | Terrabacteria group | 2,805 | 131 | 275 | 0.347 |
| *Mycobacterium smegmatis* | Bacteria | Terrabacteria group | 2,031 | 1,111 | 9,148 | 0.347 |
| *Brucella melitensis* | Bacteria | Proteobacteria | 1,287 | 684 | 1,465 | 0.346 |
| *Entamoeba histolytica* | Eukaryota | Amoebozoa | 5,656 | 641 | 1,402 | 0.344 |
| *Borrelia burgdorferi* | Bacteria | Spirochaetes | 3,168 | 186 | 361 | 0.344 |
| *Acinetobacter baumannii* | Bacteria | Proteobacteria | 3,979 | 576 | 976 | 0.344 |
| *Ureaplasma parvum* | Bacteria | Terrabacteria group | 2,244 | 113 | 275 | 0.343 |
| *Streptococcus pyogenes* | Bacteria | Terrabacteria group | 12,708 | 468 | 1,285 | 0.343 |
| *Helicobacter pylori SouthAfrica7* | Bacteria | Proteobacteria | 40,918 | 419 | 1,010 | 0.334 |
| *Fusobacterium nucleatum nucleatum* | Bacteria | Fusobacteria | 2,567 | 537 | 1,395 | 0.331 |
| *Anaplasma marginale StMaries* | Bacteria | Proteobacteria | 1,155 | 299 | 669 | 0.328 |
| *Listeria innocua* | Bacteria | Terrabacteria group | 1,164 | 659 | 1,824 | 0.318 |
| *Chlamydia trachomatis* | Bacteria | PVC group | 11,229 | 224 | 443 | 0.317 |
| *Mycoplasma putrefaciens* | Bacteria | Terrabacteria group | 15,477 | 149 | 366 | 0.315 |
| *Orientia tsutsugamushi* | Bacteria | Proteobacteria | 1,055 | 183 | 370 | 0.311 |
| *Brucella abortus* | Bacteria | Proteobacteria | 4,826 | 444 | 1,001 | 0.310 |
| *Streptococcus agalactiae* | Bacteria | Terrabacteria group | 7,335 | 268 | 353 | 0.308 |
| *Bacillus sp. 2A57CT2* | Bacteria | Terrabacteria group | 5,189 | 427 | 832 | 0.307 |
| *Staphylococcus epidermidis M23864W1* | Bacteria | Terrabacteria group | 12,643 | 460 | 696 | 0.307 |
| *Enterobacter cloacae NCTC9394* | Bacteria | Proteobacteria | 4,436 | 476 | 668 | 0.306 |
| *Streptococcus sanguinis ATCC49296* | Bacteria | Terrabacteria group | 2,991 | 342 | 559 | 0.306 |
| *Streptomyces sp. C* | Bacteria | Terrabacteria group | 1,027 | 680 | 1,348 | 0.305 |
| *Streptococcus sp. F0418* | Bacteria | Terrabacteria group | 99,893 | 192 | 203 | 0.303 |
| *Chlamydophila psittaci* | Bacteria | PVC group | 1,816 | 339 | 569 | 0.302 |
| *Propionibacterium acnes HL037PA2* | Bacteria | Terrabacteria group | 4,710 | 389 | 524 | 0.301 |
| *Vibrio harveyi* | Bacteria | Proteobacteria | 1,458 | 642 | 1,296 | 0.301 |
| *Yersinia enterocolitica* | Bacteria | Proteobacteria | 3,646 | 718 | 1,213 | 0.297 |
| *Clostridium difficile* | Bacteria | Terrabacteria group | 7,728 | 475 | 739 | 0.295 |
| *Microcystis aeruginosa* | Bacteria | Terrabacteria group | 1,451 | 418 | 799 | 0.294 |
| *Streptococcus mitis ATCC6249* | Bacteria | Terrabacteria group | 1,684 | 329 | 504 | 0.292 |
| *Paenibacillus sp. HGF7* | Bacteria | Terrabacteria group | 1,986 | 468 | 807 | 0.292 |

| | | | | | | |
|---|---|---|---|---|---|---|
| *Mycobacterium avium* | Bacteria | Terrabacteria group | 4,755 | 369 | 893 | 0.290 |
| *Mycobacterium tuberculosis H37Rv* | Bacteria | Terrabacteria group | 45,016 | 511 | 932 | 0.284 |
| *Thermotoga maritima* | Bacteria | Thermotogae | 1,108 | 288 | 316 | 0.283 |
| *Francisella tularensis tularensis* | Bacteria | Proteobacteria | 3,151 | 261 | 368 | 0.280 |
| *Sulfolobus tokodaii* | Archaea | TACK group | 2,145 | 257 | 388 | 0.279 |
| *Acinetobacter sp. ATCC27244* | Bacteria | Proteobacteria | 1,351 | 393 | 731 | 0.274 |
| *Streptococcus mitis F0392* | Bacteria | Terrabacteria group | 1,684 | 259 | 292 | 0.272 |
| *Enterobacteriaceae bacterium* | Bacteria | Proteobacteria | 374,291 | 662 | 1,121 | 0.271 |
| *Haemophilus influenzae* | Bacteria | Proteobacteria | 13,238 | 356 | 660 | 0.267 |
| *Anaerococcus tetradius* | Bacteria | Terrabacteria group | 1,334 | 244 | 267 | 0.264 |
| *Streptococcus mitis SK321* | Bacteria | Terrabacteria group | 1,684 | 215 | 215 | 0.260 |
| *Corynebacterium glutamicum* | Bacteria | Terrabacteria group | 1,299 | 519 | 615 | 0.258 |
| *Clostridium sp. 7243FAA* | Bacteria | Terrabacteria group | 1,247 | 182 | 176 | 0.255 |
| *Bartonella henselae* | Bacteria | Proteobacteria | 1,231 | 208 | 318 | 0.245 |
| *Moraxella catarrhalis* | Bacteria | Proteobacteria | 1,868 | 286 | 316 | 0.222 |

7. Minoru Kanehisa, Miho Furumichi, Mao Tanabe, Yoko Sato, and Kanae Morishima. Kegg: new perspectives on genomes, pathways, diseases and drugs. *Nucleic Acids Research*, 45(D1):D353–D361, 2016.

17. Scott Federhen. The ncbi taxonomy database. *Nucleic Acids Research*, 40(D1):D136–D143, 2011.

18. Roman L Tatusov, Darren A Natale, Igor V Garkavtsev, Tatiana A Tatusova, Uma T Shankavaram, Bachoti S Rao, Boris Kiryutin, Michael Y Galperin, Natalie D Fedorova, and Eugene V Koonin. The COG database: new developments in phylogenetic classification of proteins from complete genomes. *Nucleic Acids Research*, 29(1):22–28, 2001.

19. Michael Y Galperin, Kira S Makarova, Yuri I Wolf, and Eugene V Koonin. Expanded microbial genome coverage and improved protein family annotation in the COG database. *Nucleic Acids Research*, 43(D1):D261–D269, 2014.

20. Jaime Huerta-Cepas, Damian Szklarczyk, Kristoffer Forslund, Helen Cook, Davide Heller, Mathias C Walter, Thomas Rattei, Daniel R Mende, Shinichi Sunagawa, Michael Kuhn, et al. eggNOG 4.5: a hierarchical orthology framework with improved functional annotations for eukaryotic, prokaryotic and viral sequences. *Nucleic Acids Research*, 44(D1):D286–D293, 2015.

21. Roman L Tatusov, Eugene V Koonin, and David J Lipman. A genomic perspective on protein families. *Science*, 278(5338):631–637, 1997.

22. UniProt Consortium et al. UniProt: the universal protein knowledgebase. *Nucleic Acids Research*, 46(5):2699, 2018.

23. Daniel R Zerbino, Premanand Achuthan, Wasiu Akanni, M Ridwan Amode, Daniel Barrell, Jyothish Bhai, Konstantinos Billis, Carla Cummins, Astrid Gall, Carlos García Girón, et al. Ensembl 2018. *Nucleic Acids Research*, 46(D1):D754–D761, 2017.

24. Roland Arnold, Florian Goldenberg, Hans-Werner Mewes, and Thomas Rattei. SIMAP—the database of all-against-all protein sequence similarities and annotations with new interfaces and increased coverage. *Nucleic Acids Research*, 42(D1):D279–D284, 2013.

34. Andreas Wagner. *Robustness and Evolvability in Living Systems*. Princeton University Press, 2013.

35. Andrew L Sheldon. Equitability indices: dependence on the species count. *Ecology*, 50(3):466–467, 1969.

36. Anne MaGuarran. *Ecological diversity and its measurement*. Princeton University Press, 1988.

37. Stephen B Goodwin, LJ Spielman, JM Matuszak, SN Bergeron, and WE Fry. Clonal diversity and genetic differentiation of Phytophthora infestans populations in northern and central Mexico. *Phytopathology*, 1992.

38. Eve Runno-Paurson, Riinu Kiiker, Tiina Joutsjoki, and Asko Hannukkala. High genotypic diversity found among population of Phytophthora infestans collected in Estonia. *Fungal Biology*, 120(3):385–392, 2016.

39. Swami Iyer, Timothy Killingback, Bala Sundaram, and Zhen Wang. Attack robustness and centrality of complex networks. *PLoS One*, 8(4):e59613, 2013.

40. Marcel Salathé, Maria Kazandjieva, Jung Woo Lee, Philip Levis, Marcus W Feldman, and James H Jones. A high-resolution human contact network for infectious disease transmission. *Proceedings of the National Academy of Sciences*, page 201009094, 2010.

41. Miroslav Fiedler. Algebraic connectivity of graphs. *Czechoslovak Mathematical Journal*, 23(2):298–305, 1973.

42. Michael Begon, John L Harper, Colin R Townsend, et al. *Ecology. Individuals, populations and communities.* Blackwell Scientific Publications, 1986.

43. S Mohan Jain and Subhash C Minocha. *Molecular biology of woody plants*, volume 1. Springer Science & Business Media, 2013.

49. Ronald S Burt. *Structural holes: The social structure of competition.* Harvard University Press, 2009.

56. Béla Bollobás. *Modern graph theory*, volume 184. Springer Science & Business Media, 2013.

65. David Easley and Jon Kleinberg. *Networks, crowds, and markets: Reasoning about a highly connected world*. Cambridge University Press, 2010.

66. Robert R Sokal. Biometry: the principles and practice of statistics. *Biological Research*, 1995.

67. HT Reynolds. Nonparametric partial correlation and causal analysis. *Sociological Methods & Research*, 2(3):376–392, 1974.

68. Dana Quade. Nonparametric partial correlation. *Measurement in the Social Sciences*, pages 369–398, 1974.

69. William Jay Conover and William Jay Conover. Practical nonparametric statistics. 1980.

75. Helen Berman, Kim Henrick, Haruki Nakamura, and John L Markley. The worldwide protein data bank (wwpdb): ensuring a single, uniform archive of pdb data. *Nucleic Acids Research*, 35(suppl_1):D301–D303, 2006.

82. Hao Luo, Yan Lin, Feng Gao, Chun-Ting Zhang, and Ren Zhang. DEG 10, an update of the database of essential genes that includes both protein-coding genes and noncoding genomic elements. *Nucleic Acids Research*, 42(D1):D574–D580, 2013.
